# Development of 3-in-1 nanotherapeutic strategies for ovarian cancer

**DOI:** 10.1101/2024.07.17.604002

**Authors:** Emma Durocher, Sean McGrath, Esha Gahunia, Naomi Matsuura, Suresh Gadde

**Author notes:** Corresponding author: Suresh Gadde.

## Abstract

**TOC.**
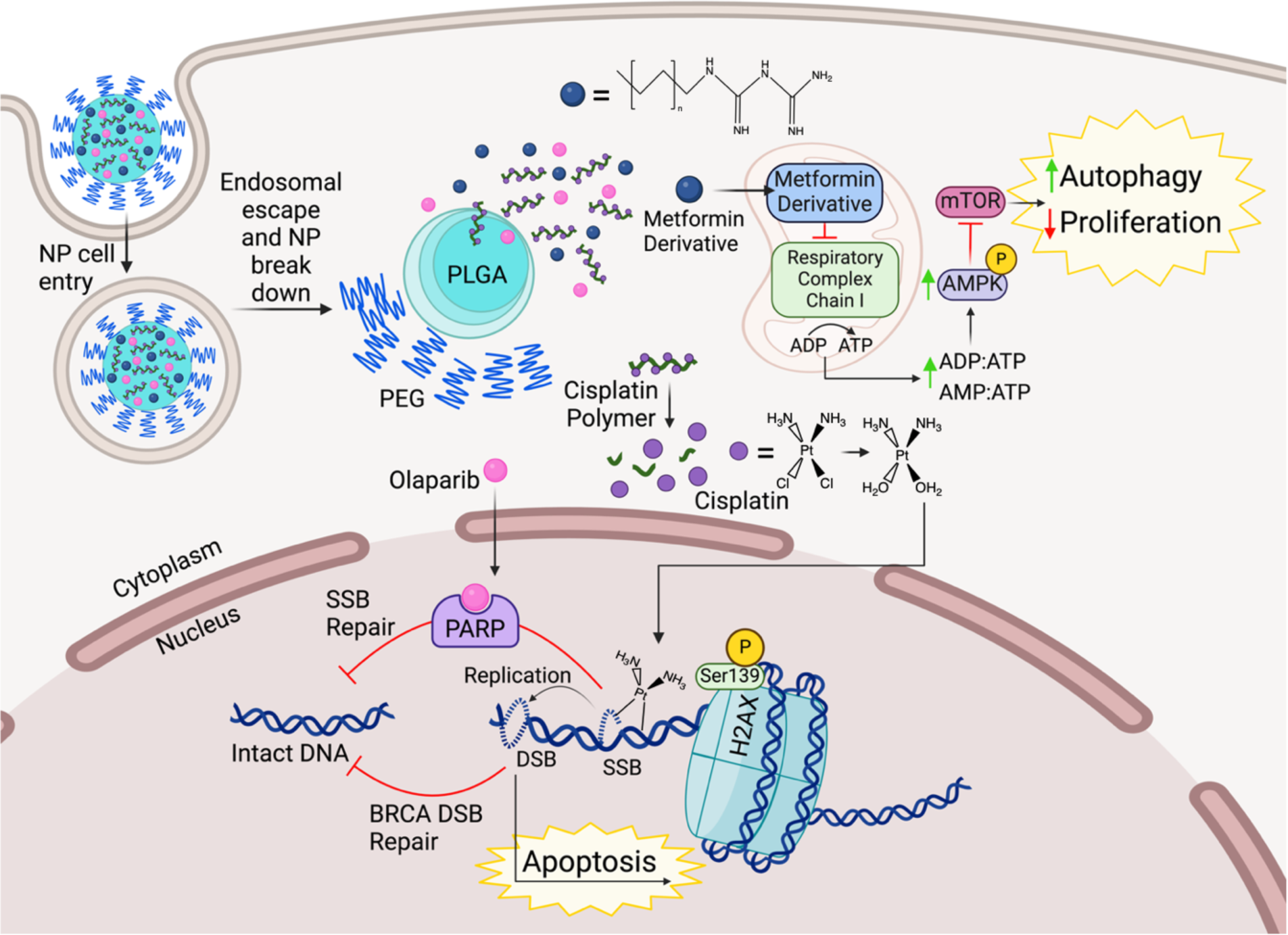

Among gynecological cancers, ovarian cancer causes the most fatality. Platin-based chemotherapy is the primary therapeutic option, but it is limited by a variety of drug resistance mechanisms. Ovarian cancer is a complex and challenging disease to treat, and combination approaches have shown stronger efficacy than a single drug alone. However, they still need to overcome challenges, such as the non-selective distribution of drugs, and side effects caused by each drug in the combination. To overcome these issues, here we explored a 3-in-1 combination nanotherapeutic approach containing cisplatin, olaparib, and metformin for ovarian cancer. To encapsulate hydrophilic cisplatin and metformin inside the nanoparticle (NP) core, we developed cisplatin polymer prodrugs and metformin derivatives. Our results showed successful development of 3-in-1 NPs containing cisplatin, olaparib, and metformin, and they are stable in the physiological conditions. In vitro evaluation showed each agent in the 3-in-1 NPs is active and exerts therapeutic effects, contributing to ovarian cancer cell killing at lower concentrations. These results provide insight into developing novel nanotherapeutic strategies for improving ovarian cancer treatment.

## INTRODUCTION

Among gynecological cancers, ovarian cancer is the leading cause of death in women^1^. Currently, platinum-based chemotherapy is the standard treatment regimen for patients. Of the platinum-based chemotherapeutic options, cisplatin is a potent and effective cancer cell killing agent (Scheme 1).^1^ However, cisplatin efficacy remains limited due to side effects caused by the systemic, non-selective distribution of the drug and the development of resistance to platin. Among different resistance mechanisms that occur before or after cisplatin-DNA adduct formation,^2^ Poly (ADP-ribose) polymerase (PARP)-mediated DNA repair is a predominant mechanism associated with developing cisplatin resistance in ovarian cancer (Scheme 1).^3^ PARP works to detect single-strand breaks in DNA via its zinc-finger domain and activates the DNA repair proteins.^3, 4^ Additionally, mTOR has been shown to overexpress in advanced-stage ovarian cancer tumors, which also prevents cisplatin-induced apoptosis via positively regulating tumor proliferation and cell cycle progression (Scheme 1).^5, 6^ In addition to platinum resistance, systemic non-selective distribution causes low tolerability and severe toxicities.^7^ Patients on long-term cisplatin use have reported side effects, including but not limited to vomiting, nausea, nephrotoxicity, and myelosuppression, as well as damage to healthy tissue.^7^

Ovarian cancer is a complex and challenging disease to treat, and combination therapeutic approaches have shown to have better efficacy with decreased resistance development and increased overall survival than single drug therapies.^8–10^ In this context, cisplatin, in combination with olaparib, a PARP inhibitor, was explored in several preclinical and clinical settings. Due to favorable results, olaparib has been added to the treatment regimen as a maintenance therapy for ovarian cancer patients.^8–10^ Furthermore, metformin is a clinically approved antidiabetic drug that has been recently reported to have anticancer effects. Several studies have used metformin to sensitize ovarian cancer cells to cisplatin by indirectly increasing AMPK activation.^11, 12^ AMPK is a negative regulator of mTOR, and since inhibiting mTOR has proven to be difficult, targeting AMPK, a protein upstream, has shown to provide better autophagic outcomes.^13, 14^ While these combination therapies are effective, they still need to overcome multiple drawbacks to reach their full potential, such as having synergistic drug ratios in tumors to achieve maximum benefits, minimizing toxic side effects, and overcoming the dosing schedule.^8,15^

**Scheme 1.**
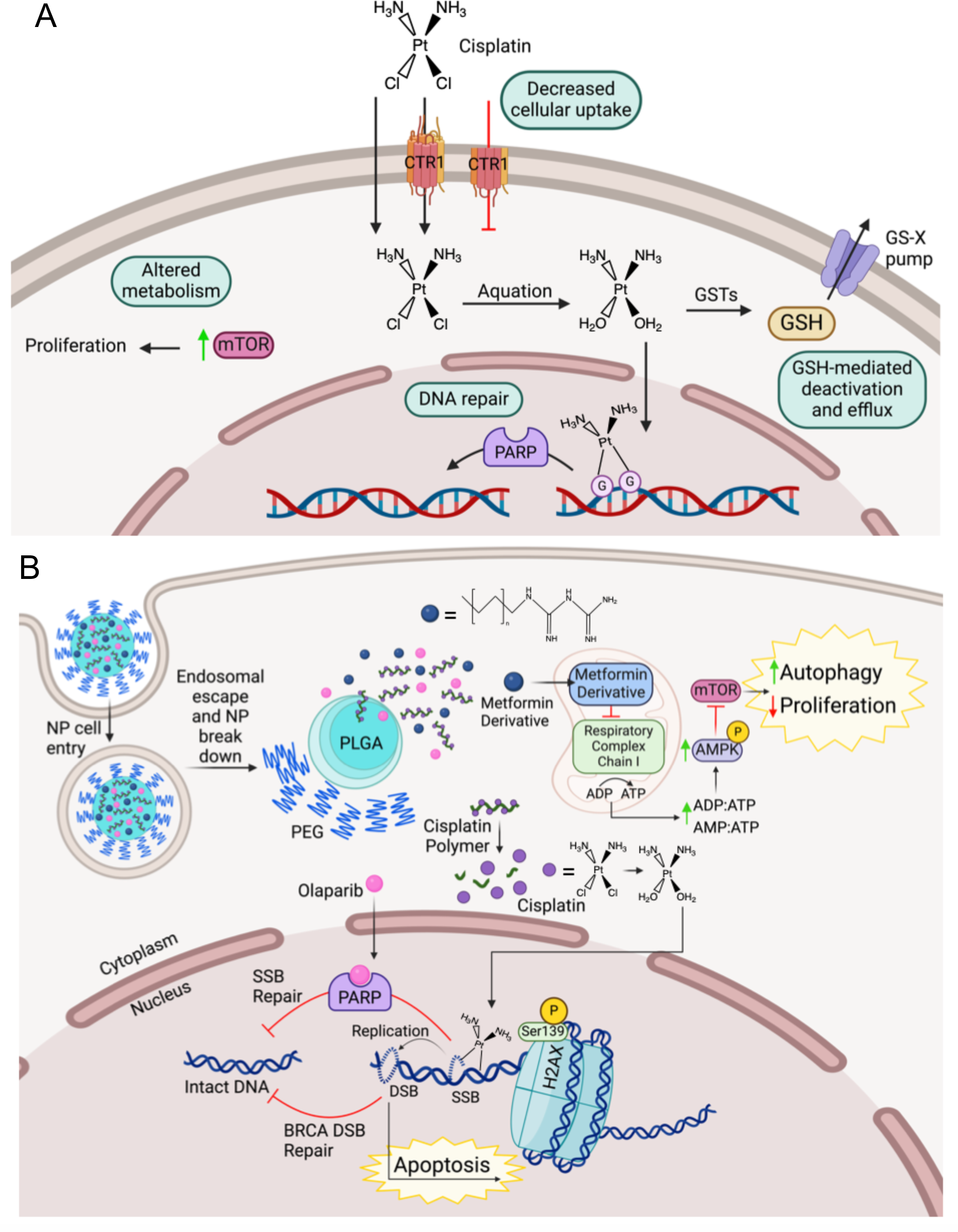
Cisplatin mechanism of action and resistance pathways addressed by 3-in-1 NP treatment. (A) Cisplatin enters the cells either by passive diffusion or through the CTR1 transporter. Once inside, cisplatin undergoes aquation and crosses into the nucleus where it forms adducts with DNA. Resistance mechanisms are labelled 1-4. (1) Excision of the N-terminal domain on CTR1 prevents facilitated cisplatin uptake. (2) GSTs catalyze the conjugation of GSH to active cisplatin, which promotes efflux via GS-X pumps. (3) DNA repair of cisplatin DNA adducts prevent cell death. (4) Upregulated mTOR levels confers an altered metabolism that promotes proliferation. (B) 3-in-1 NP enters through the endocytic pathway before undergoing endosomal escape and breakdown of the PLGA-PEG polymers, releasing the contents. The metformin derivative (blue circle) inhibits respiratory complex chain I in the mitochondria, thereby increasing the ratio of ADP and AMP to ATP, leading to AMPK activation, inhibition of mTOR. Cisplatin polymer is reduced releasing cisplatin (purple circle). Cisplatin undergoes aquation, before crossing into the nucleus to form DNA adducts. Ser139 on H2AX becomes phosphorylated following adjacent DNA damage. Olaparib (pink circle) binds to and cleaves PARP preventing SSB repair of cisplatin induced DNA damage. PARP inhibition coupled with mutated BRCA DSB repair causes synthetic lethality, leading to apoptosis.

Nanoparticles (NPs) are nanometer-sized particles that can encapsulate single or multiple drugs with different physicochemical properties, protect them from degradation, and deliver them to specific organs and tissues.^16^ In the case of cancer nanomedicines, NPs can selectively accumulate in tumor sites via active or passive targeting mechanisms, such as the enhanced permeation and retention and newly proposed alternative mechanisms.^16, 17^ For cancer combination therapies, NPs’ have a multitude of advantages. They can encapsulate combination drugs with different physicochemical properties in synergistic drug ratios, deliver them to tumors to induce overlap in pharmacological profiles, maximize the therapeutic effects in the tumors, and minimize the side effects caused by each drug in the combination.^16^

Since drug resistance is a major issue in developing potent therapies for ovarian cancer, we rationalized that triple drug combination therapies are effective ways to combat the disease. To this end, we explored combination therapeutic strategies with cisplatin, olaparib, and metformin to synergistically improve and enhance cisplatin cytotoxicity while blocking multiple drug-resistant mechanisms. We developed 3-in-1 NP systems containing cisplatin, olaparib, and metformin to induce overlap in their pharmacokinetic profiles and synergistic therapeutic effects (Scheme 1). For encapsulating 3 drugs in optimal ratios in 3-in-1 NPs, we employed polymer-prodrug strategies to develop cisplatin-based polymer prodrugs (DTB and AA polymer) and metformin derivatives. Upon synthesis and characterization of these prodrugs/derivatives, we utilized them to develop 3-in-1 NPs and studied their effects using in vitro ovarian cancer models. Our studies showed that 3-in-1 NPs are effective in encapsulating all 3 agents inside the NPs and are effective in inducing their respective therapeutic effects.

## RESULTS

### Development of cisplatin-based polymer prodrugs (DTB and AA), and metformin derivatives (C_16_-Met and C_6_-Met)

Poly lactic-co-glycolic acid-polyethylene glycol (PLGA-PEG) based polymeric NPs platforms have several advantages, such as stability, scalability, biodegradability, and easy to manipulate.^18, 19^ Additionally, these systems are easy to modify to accommodate multiple drugs with different physicochemical properties for multidrug delivery. Previous studies successfully developed multidrug delivery platforms by blending PLGA-PEG polymers with different polymer-drug conjugates, where drugs are covalently conjugated to side chains of polymers.^20–22^ In this study, to develop 3-in-1 NP platforms containing synergistic ratios of cisplatin, olaparib, and metformin, we first opted for polymer prodrugs to overcome inherent limitations. In order to blend with PLGA-PEG polymers, we developed two cisplatin polymer-prodrugs to increase hydrophobicity without altering the pharmacological activity of cisplatin, and to encapsulate inside the NPs’ hydrophobic core. For this, hydroxyl groups on Oxoplatin (**Fig. 2A**) were reacted with two different dicarboxylic acids to produce DTB and AA polymers. DTB Polymer contains a disulfide bond in the linker moiety connecting cisplatin units together for GSH scavenging potential (**Fig. 2A**). AA Polymer was created to serve as a control for DTB Polymer, thereby mimicking the structure, except lacking the disulfide bond (**Fig. 2A**). Next, similar to cisplatin, metformin, another drug in our combination, is also hydrophilic and orally administrated for diabetic treatments.^11^ However, to produce synergistic effects, we reasoned that all 3 drugs need to be in the tumor microenvironment to induce pharmacological overlap. To improve metformin loading into PLGA-PEG NPs, two different metformin derivatives were synthesized by reacting dicyandiamide with 2 different long chain amines to create C_16_-Met and C_6_-Met.^23^ They have varying hydrocarbon chain lengths to modulate hydrophobicity and encapsulation (**Fig. 2B**). All structures were confirmed by ^1^H NMR and/or ^13^C NMR (**Fig. 2, Fig. S1-5**). The percent of platinum in DTB and AA polymers was measured by ICP-MS.

**Figure 1.**
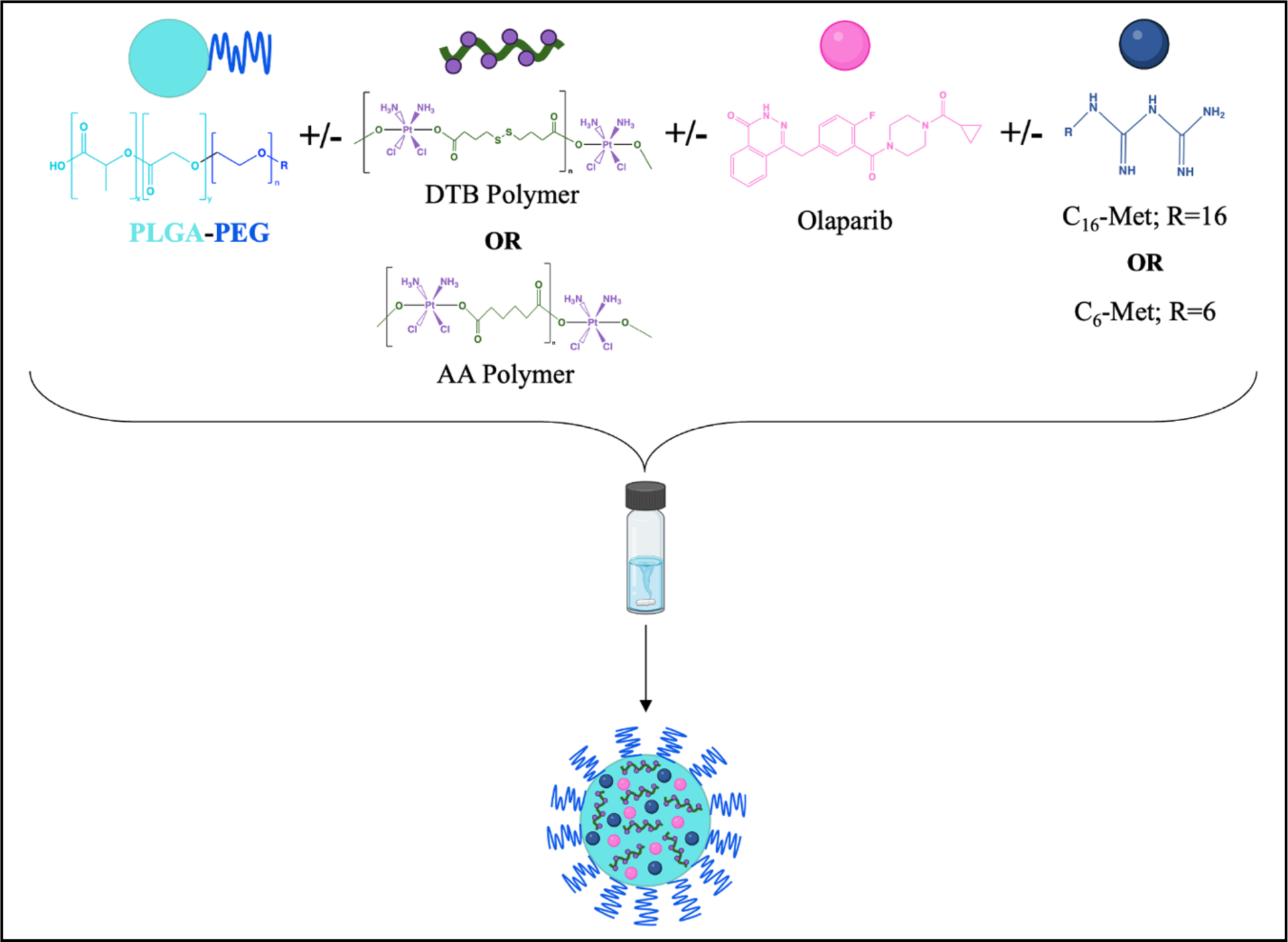
Development of 3-in-1 NPs containing cisplatin, olaparib, and metformin.

**Figure 2.**
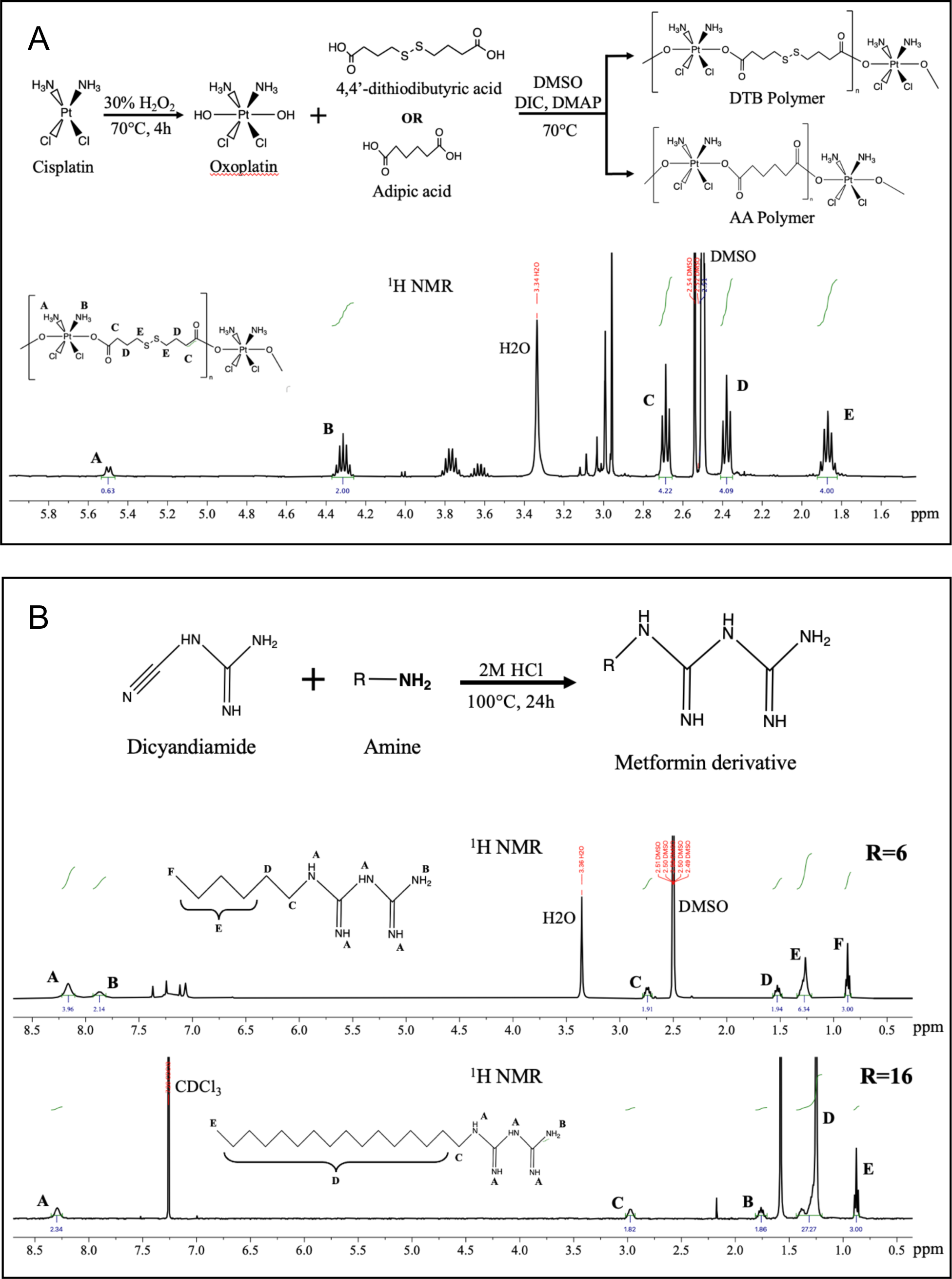
Synthesis and characterization of cisplatin polymer prodrugs and metformin derivatives. (A) Synthetic scheme of DTB and AA Polymer (top). Cisplatin reacted with 30% hydrogen peroxide (H_2_O_2_) at 70°C for 4 hours to form oxoplatin. Oxoplatin reacted with wither 4,4’-dithiodibutyric acid or adipic acid in DMSO with DIC and DMAP activators at 70°C to form DTB polymer or AA Polymer, respectively. ^1^H NMR spectrum of DTB Polymer in DMSO-d6 (bottom). (B) Synthetic scheme of metformin derivatives (top). Dicyandiamide reacted with an amine compound (R=hydrocarbon chain) in 2M HCl at 100°C for 24 hours to form the metformin derivatives, C_16_-Met and C_6_-Met. ^1^H NMR spectrum of C_6_-Met (middle) and C_16_-Met (bottom) in DMSO-d6 and CDCl_3_, respectively. Peaks corresponding to specific hydrogens in the compound are identified using alphabetical labeling for all NMR spectra.

### Cisplatin, olaparib, and metformin combinations display synergy in SKOV-3 cells

Next, we employed MTT assays to determine synergistic ratios of the drug combinations to encapsulate them in our 3-in-1 NP system. Our results showed most combination treatments resulted in lower cell viability compared to either single agent (**Fig. 3A**). The data presented in **fig. 3A**, along with MTT data of cisplatin and metformin, and olaparib and metformin combinations in SKOV-3 cells were inputted into either Synergy Finder (**Fig. 3B**) or CompuSyn software (**Fig. 3C**) to determine the synergistic ratio between the dual drug combinations. Synergy Finder uses the HSA reference method to calculate synergy, in which a positive synergy score indicates a synergistic relationship.^24^ The identified concentration ranges that demonstrated synergy in combination were at the lower end, indicating a synergistic relationship can be achieved while lowering the concentration of each drug (**Fig. 3B**). CompuSyn uses the established Chou-Talalay method for synergy calculation, in which combination index (CI) values < 1 indicate synergy.^25^ The concentration ranges corresponding to CI<1 corroborate the data obtained from Synergy Finder (**Fig. 3C**). Lastly, a final synergy study was conducted to determine the synergistic relationship between all three cisplatin, olaparib, and metformin in combination, revealing concentration ranges corresponding to CI<1 calculated by CompuSyn software (**Fig. 3C**).

**Figure 3.**
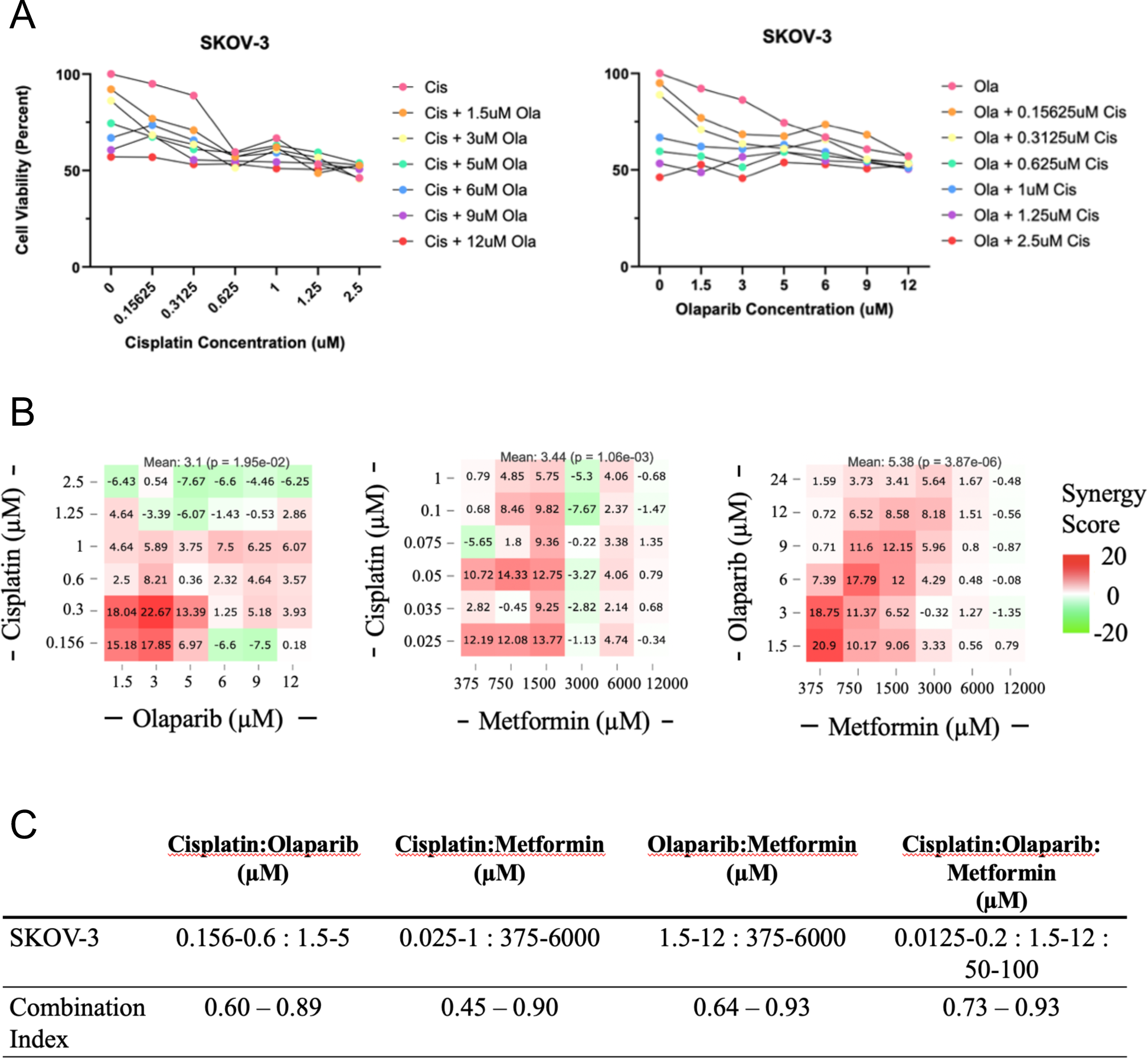
Synergistic ratios between cisplatin, olaparib, and/or metformin in SKOV-3 cells. (A) SKOV-3 cell viability following treatment with cisplatin, olaparib and the two agents in combination for 72 hours plotted either against cisplatin concentration (left) or olaparib concentration (right). Data measured by MTT assays (n=1). MTT data was inputted into either (B) Synergy Finder or (C) CompuSyn to obtain synergy scores and combination index (CI) values, respectively for cisplatin and olaparib, cisplatin and metformin, and olaparib and metformin combinations in SKOV-3 cells. Cisplatin, olaparib, and metformin combination CI values are shown in (C) (n=1). The synergy scores were calculated using the HSA reference model. CompuSyn calculates CI values using the Chou Talalay method.

### Development of 3-in-1 PLGA-PEG NPs containing cisplatin, olaparib and metformin

We next developed 3-in-1 NPs and appropriate control NPs using the nanoprecipitation method by blending different ratios of polymers, polymer prodrugs, and/or metformin derivatives and/or olaparib (**Fig. 4A**).^26–28^ The various NPs synthesized include E-NP (Empty-NP), single agent AA-NP, DTB-NP, Ola-NP, C_16_-Met-NP, C_6_-Met-NP, and 3-in-1 DTB-Ola-C_16_-Met-NP, and DTB-Ola-C_6_-Met-NP. Particle size (nm) and polydispersity index (PDI) of all NPs were measured using dynamic light scattering (**Fig. 4B and C, Fig. S10**). The mean size and PDI between all NPs were consistent, ranging between 85.4-125.6 nm, and 0.069-0.11, respectively. NPs size did not change considerably going from single drug-NPs to 3-in-1 NPs. The uniform spherical morphology of DTB-Ola-C_16_-Met-NP (representative for all NPs) was confirmed by transmission electron microscopy (TEM) images at 84.2k magnification (**Fig. 4D**). The surface potential of AA-NP, DTB-NP, DTB-Ola-C_16_-Met-NP, and E-NP were also determined using dynamic light scattering (**Fig. 4E**). DTB-NP revealed a negative surface charge of −7.96±2.24 mV, while DTB-Ola-C_16_-Met-NP displayed a near-neutral charge of 0.51±0.25 mV. Moreover, cisplatin encapsulation efficiency (EE) was measured by ICP-MS. DTB-Ola-C_16_-Met-NP was shown to have the highest cisplatin EE of 30.31±3.54%, notably higher than DTB-Ola-C_6_-Met-NP EE of 12.01% (n=1).

**Figure 4.**
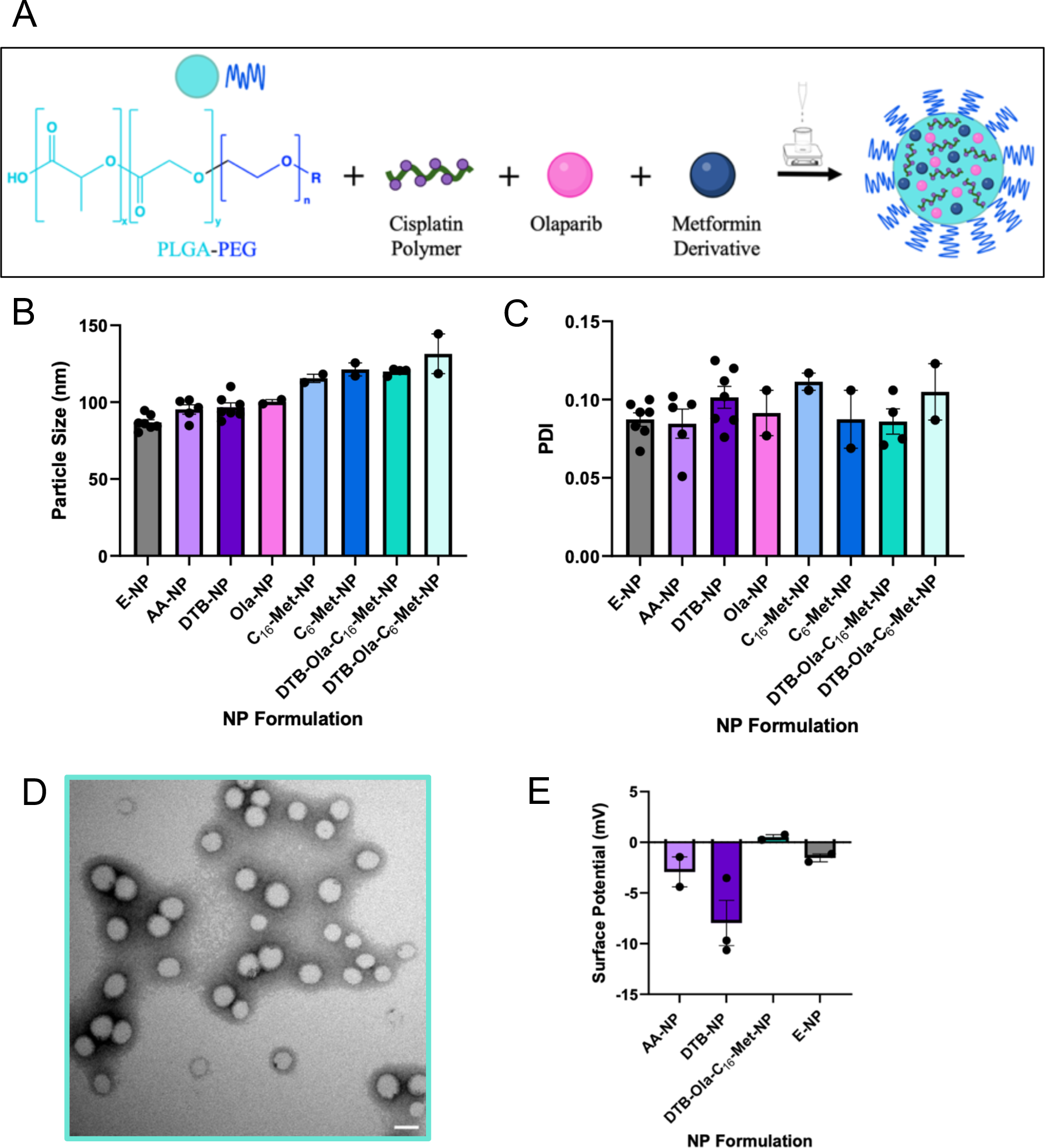
Synthesis and characterization of cisplatin-olaparib-metformin NPs. (A) Synthetic scheme of cisplatin polymer, olaparib, and metformin derivative loaded PLGA-PEG NPs by nanoprecipitation. (B) Particle size (nm) and (C) PDI of empty (E), single, and triple drug NP formulations measured by dynamic light scattering (n=7,5,4, or 2). (D) TEM image of DTB-Ola-C_16_-Met-NP at 84.2k magnification. Scale bar: 50nm. (E) Surface potential (mV) of AA-, DTB-, DTB-Ola-C_16_-Met-, and E-NP determined by dynamic light scattering (n=3 or 2). Data presented as means ± SEM.

### 3-in-1 NPs and E-NPs are stable across temperature and FBS conditions

To assess the stability of our 3-in-1 DTB-Ola-C_16_-Met-NP and E-NP at 4°C, size and PDI measurements were made immediately after synthesis and again after 60 days of storage (**Fig. 5A**).^28^ Data revealed no significant change of either size or PDI, indicating NPs are stable for long term. Additionally, the serum stability of NPs was assessed by incubating them in 0, 5, or 10% FBS at both 37°C (**Fig. 5B**) and 25°C (**Fig. 5C**) for different time periods.^28, 29^ Across the 24-hour time interval within each condition, there was no observable change in size (**Fig. 5B and C, left**) or PDI (**Fig. 5B and C, right**). As the percent FBS increased, size trended downwards, whereas PDI gradually increased. Both NPs displayed the same stability trends irrespective of the starting size. Between temperature settings, PDI showed no difference, whereas NP size at each condition was slightly greater at 37°C compared to measurements taken at 25°C. Taken altogether, both NPs showed negligible changes in size across percent FBS, time, and temperature, demonstrating size stability in FBS across a period of 24 hours.

**Figure 5.**
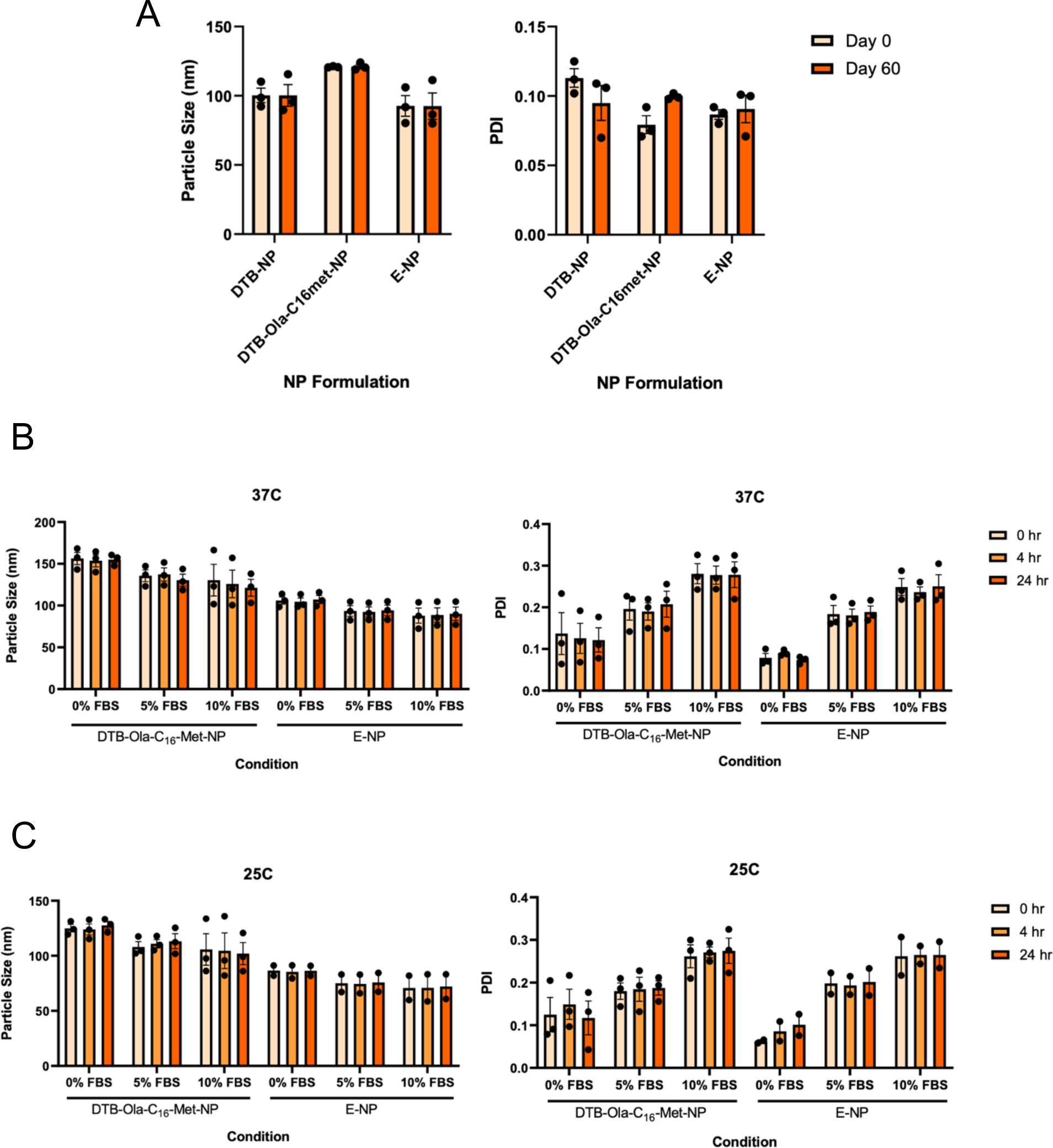
Size and PDI stability of DTB-Ola-C_16_-Met- and E-NP. (A) Particle size (nm) and PDI of DTB-, DTB-Ola-C_16_-Met-, and E-NP measured by dynamic light scattering immediately after synthesis (Day 0) and again after 60 days of storage at 4°C. Data is presented as means ± SEM (n=3). (B, C) Particle size (nm) and PDI stability of DTB-Ola-C_16_-Met- and E-NP following incubation at either (B) 37°C or (C) 25°C in 0, 5, and 10% FBS measured at 0, 4, and 24 hours. Data is presented as means ± SEM (n=3 or 2).

### Validation of individual drug pharmacological effects within the 3-in-1 NP, and its capacity to induce toxicity in ovarian cancer cells

Before assessing the therapeutic effects of 3-in-1 NPs, we first evaluated each agent in the 3-in-1 NP for their contribution to the overall therapeutic effect. Cisplatin induces cytotoxicity via DNA damage by forming DNA adducts.^2^ For this, we evaluated the capacity of AA and DTB Polymers in NP systems to undergo Pt(IV) to Pt(II) reduction and form DNA adducts to induce DNA damage. To this end, we performed γH2AX assays, in which upon DNA damage, the histone, H2AX, becomes phosphorylated (referred to as γH2AX) and acts as an early biomarker of apoptosis.^30^ Here, we measured γH2AX levels by flow cytometry. SKOV-3 cells were treated with DTB-NP, AA-NP, DTB-Ola-C_16_-Met-NP, and E-NP for 24 hours and assessed for DNA damage using flow cytometry. We observed a similar increase in median fluorescence intensity of the second peak following both AA- and DTB-NP treatment, compared to E-NP (**Fig. S11**). The second peak is believed to be caused by pronounced DNA damage induced by cisplatin, whereas the first observable peak is suggested to correspond to physiological DNA damage levels. Additionally, there was a greater observed shift in the second peak toward higher intensity following DTB-Ola-C_16_-Met-NP treatment (**Fig. 6A**), showing cisplatin prodrug polymers in 3-in-1 NPs are effective and induce DNA damage. Additionally, it is important to note that cisplatin concentration in the 3-in-1 NPs (0.25uM) was 32-fold lower than that of AA- and DTB-NP and the cisplatin alone control (8uM).

**Figure 6.**
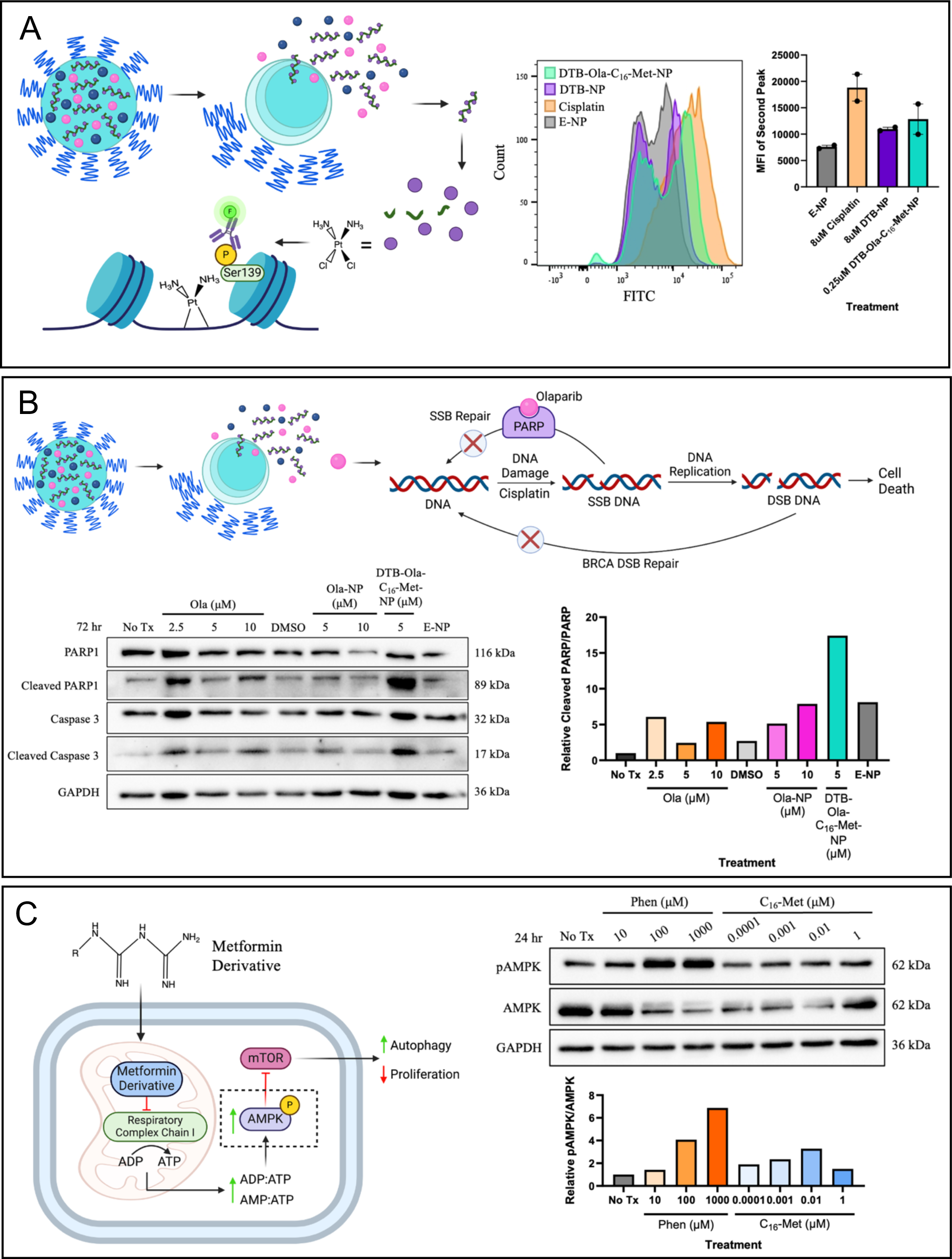
Validation of individual drug pharmacological effects within the triple drug NP. (A) Schematic representation of DTB-Ola-C_16_-Met-NP and DTB polymer break down, releasing cisplatin, which can then form DNA adducts. Ser139 on Histone 2AX becomes phosphorylated following adjacent DNA damage which is tagged with a FITC-IgG antibody. Representative flow cytometry histogram of FITC-tagged γH2AX induced by cisplatin in SKOV-3 cells (left) and median fluorescence intensity (MFI, right) following E-NP, 8μM cisplatin, 8μM DTB-NP or 0.25μM DTB-Ola-C_16_-Met-NP treatment for 24 hours. Data is presented as means ± SEM (n=2). (B) Schematic representation of DTB-Ola-C_16_-Met-NP break down, releasing olaparib to undergo its mechanism of action. Olaparib inhibits PARP preventing SSB repair of cisplatin induced DNA damage. PARP inhibition coupled with mutated BRCA DSB repair causes synthetic lethality, leading to cell death. SSB, single strand break; DSB, double strand break. Western blot of PARP1 and caspase 3 and their cleaved counterparts in SKOV-3 cells after Ola-NP, DTB-Ola-C_16_-Met-NP, and E-NP treatment for 72 hours. Relative cleaved PARP1/PARP1 ratio quantified using Fiji. Data normalized to No Tx control, which is set to 1 (n=1). (C) Schematic representation of either free C_16_-Met or C_6_-Met derivative proposed mechanism of action. The metformin derivative inhibits respiratory complex chain I, thereby increasing the ratio of ADP and AMP to ATP, leading to AMPK activation, inhibition of mTOR and ultimately increased autophagy and decreased cell proliferation. The dotted box shows the step in the pathway for which has been blotted. Western blot of phosphorylated AMPK (pAMPK) in SKOV-3 cells after C_16_-Met treatment for 24 hours. Phenformin was used as a positive control. Relative pAMPK/AMPK ratio quantified using Fiji. Data normalized to No Tx control, which is set to 1 (n=1).

**Figure 7.**
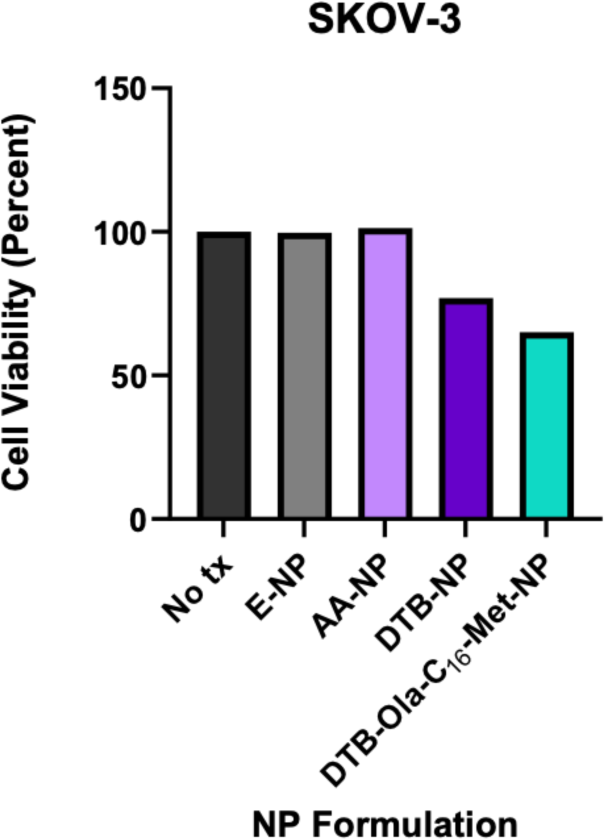
Cytotoxicity of AA-, DTB- and DTB-Ola-C_16_-Met-NPs in SKOV-3 cells. (C) SKOV-3 cell viability following treatment with E-NP, AA-NP, DTB-NP, and DTB-Ola-C_16_-Met-NP for 72 hours, with dosing every 24 hours (n=1). Cisplatin concentration in each treatment was 100nM. Viability was measured by MTT assays. Treatments were normalized to No tx (treatment) control cells.

Next, we validated the pharmacological activity of olaparib in 3-in-1 NPs by treating SKOV-3 cells with olaparib, Ola-NP, DTB-Ola-C_16_-Met-NP, and E-NP for 72 hours. We analyzed cleaved PARP and caspase-3 expression by western blot analysis (**Fig. 6B**). Our results showed PARP and caspase-3 cleaved products are more abundant following DTB-Ola-C_16_-Met-NP treatment compared to Ola- and E-NP suggesting olaparib in our 3-in-1 NPs is active and inhibits PARP mediated DNA repair. For the third agent in our 3-in-1 NP, metformin derivative, we analyzed its target, AMPK activation via WB. Unlike previous experiments, here treated SKOV-3 cells with free C_16_-Met and C_6_-Met (0.0001-1μM) for 24 hours, but not 3-in-1 NPs due to challenges in experimental conditions to observe AMPK activation. We observed a concentration dependent increase in pAMPK expression induced by both derivatives (**Fig. 6C and S13**). However, a drop in pAMPK was noticed at the highest concentration (1μM).

Following the confirmation that, all drugs in the 3-in-1 NPs are active, we performed preliminary cytotoxicity assays of our 3-in-1 NPs. SKOV-3 cells were treated with AA-NP, DTB-NP, and DTB-Ola-C_16_-Met-NP for 72 hours, and the cytotoxicity was analyzed using MTT assays. While our results show considerable cell death induced by 3-in-1 NPs when compared to control NPs, we did not observe potent cytotoxicity. This might be due to the fact that the drug concentrations in our NPs are several folds lower than the cytotoxicity observed by free drug concentrations.

## DISCUSSION

Ovarian cancer is the most fatal gynecological cancer among women.^1^ However, conventional platinum-based chemotherapy limits treatment effectiveness and tolerability due to resistance development and systemic distribution.^1, 2^ Previous studies have employed combination therapy approaches to overcome cisplatin resistance.^8, 15^ Particularly, olaparib, a small molecule drug used to inhibit PARP-mediated DNA repair, has improved cisplatin sensitivity, and is therefore used as a maintenance therapy in the clinic.^31^ Additionally, metformin has garnered attention for its anticancer effects by indirectly inhibiting mTOR, thereby altering metabolism toward a catabolic state to ultimately cause autophagy.^11, 12^ Given the complexity of ovarian cancer, targeting multiple pathways may provide longer lasting remission. Improvements to combination therapy using NP delivery systems have also been investigated to prevent off target effects, thereby increasing drug payload to the tumor. Although several studies have explored NP delivery of cisplatin, olaparib or metformin, NP development encapsulating a triple drug combination for ovarian cancer treatment has yet to be explored.^32–35^ Here, we sought to develop a novel NP-based therapy encapsulating a cisplatin polymer prodrug, olaparib, and a metformin derivative in a single nanocarrier for a triple therapeutic effect for ovarian cancer therapy.

Our results showed successful encapsulation of all 3 agents in a single NP, and these NPs have ideal sizes and surface charges and are uniform and stable. Additionally, flow cytometry and WB analysis showed that 3-in-1 NPs can induce DNA damage and PARP inhibition to produce synergistic cytotoxic effects. Also, it is important to note that 3-in-1 NPs are able to exert better effects at lower concentrations than the drugs alone, as cisplatin and olaparib concentrations in 3-in-1 NPs are lower than in free drug treatments (Fig. 6A and B). A similar trend was observed with metformin, as metformin derivatives showed better AMPK activation than the control metformin. Overall, our results showed agents in 3-in-1 NPs are effective and can work together to induce synergistic effects.

While these results are promising, further optimization of 3-in-1 NPs is needed to achieve maximum therapeutic benefits. While previous studies showed including S-S bonds in the polymers/NP platforms enables GSH scavenging/responsive cisplatin release and minimizes deactivation and efflux of cisplatin,^36–38^ in our experimental conditions we did not observe this effect with our 3-in-1 NPs, which also contain S-S bonds via DTB polymer (data not shown). Since PLGA-PEG and DTB polymers are blended to develop 3-in-1 NPs, PLGA hydrophobic core might have limited GSH responsive/scavenging ability of DTB polymer. In our 3-in-1 NPs we have used 50K MW PLGA, and lowering PLGA MW and modulating distance between Platinum to S-S moiety in DTB polymers might help to improve the GSH responsive/scavenging ability of our 3-in-1 NPs.

While our preliminary cytotoxicity results show 3-in-1 NPs are effective, in observed treatment conditions, they did not induce more than 50% cytotoxicity. It is important to note that cisplatin concentrations used in these studies (100nM) are greatly below reported IC50 values.^39, 40^ The 100nM cisplatin concentration was chosen based on the synergy findings. Additionally, PLGA-PEG NPs have controlled drug release kinetics, and we employed 50K MW PLGA for our 3-in-1 NPs. Therefore, it is possible that 72-hour time frame used for all cytotoxicity studies may not be sufficient to observe substantial cell death. Our DNA damage, PARP cleavage, and pAMPK expression data does provide supporting evidence that cytotoxicity from DTB-Ola-C_16_-Met-NP should be attainable; however, this may only be possible with longer treatment times.

## CONCLUSION

Ovarian cancer is the most lethal gynecological cancer^1^. However, response to treatment remains limited due to chemotherapy resistance and systemic toxicities. Here, we developed a novel 3-in-1 nanotherapeutic strategy using cisplatin, olaparib, and metformin. We successfully synthesized cisplatin polymer prodrugs and metformin derivatives to ensure adequate encapsulation inside the PLGA-PEG NP hydrophobic core. Our synergy findings demonstrated that all three agents operate synergistically when combined at certain concentrations. PLGA-PEG NPs encapsulating AA or DTB Polymer, olaparib, and/or C_16_-Met or C_6_-Met were formulated while considering the identified synergistic ratio. Final formulations were optimized according to morphology, size, PDI, surface charge, and EE and tested for their therapeutic effects. DNA damage induced by cisplatin from encapsulated AA or DTB Polymer was shown by using γH2AX flow cytometry assays. Additionally, our studies showed preliminary evidence of PARP and caspase-3 cleavage as well as pAMPK expression caused by olaparib and free C_16_-Met/C_6_-Met, respectively, by western blot analysis. These findings can lead to the development of potent combination nanotherapeutic approaches for ovarian cancer treatment.

## MATERIALS AND METHODS

### Materials

Cisplatin (Cis) was purchased from Sigma Aldrich (Cat # 479306-5G). Olaparib (Ola), and metformin (Met) were purchased from Selleckchem (Cat # S1060, # S1950, # S1208, respectively).

### Synthesis and characterization of polymer prodrugs and derivatives

All synthesis products were characterized by either nuclear magnetic resonance (NMR) using a Bruker AVANCE II 400, inductively coupled plasma mass spectrometry (ICP-MS), and/or electrospray ionisation mass spectrometry (ESI-MS).

### Oxoplatin [PtCl_2_(NH_3_)_2_OH_2_]

Cisplatin (496.2mg, 1.65 mmol) was added to a 50mL round bottom (RB) flask with water (3mL). 30% hydrogen peroxide (16mL, BioShop) was added, the temperature was set at 75°C, and the reaction proceeded for 4 hours.^30^ After cooling, the reaction mixture was filtered through 5μm filter paper. The product was then washed separately with water and diethyl ether and then left to dry again for roughly 1 hour after each wash. The product was collected in a 20mL glass vial and placed on the vacuum for 3 days. Oxoplatin was collected in 49.6% (273.6mg) yield.

### DTB Polymer

4,4’-dithiodibutyric acid (35.56mg, 0.15mmol, Sigma Aldrich, Cat # C15605-10G) in DMSO (1mL) was added to a 20mL vial. N,N’-diisopropylcarbodiimide (DIC, 5uL, 3 equiv., Alfa Aesar, Cat # A19292) and 4-dimethylaminopyridine (DMAP, 54.68mg, 3 equiv., Alfa Aesar, Cat # A13016) were added as catalysts for esterification. Oxoplatin (50mg, 1 equiv.) in DMSO was added to the reaction. The reaction mixture was heated to 70°C and proceeded overnight. After cooling the reaction mixture, the solution was lyophilized, and the resulting product was dialyzed in methanol using SnakeSkin dialysis tubing (3500 Da MWCO, Thermo Fisher, Cat # 68035) for 24 hours. The product was collected via rotary evaporation, followed by lyophilization. Product was stored at 4°C. ^1^H NMR (400 MHz, DMSO-d6): δ 5.46 (d, NH3), 4.27 (hept, NH3), 2.65 (t, CH2), 2.34 (t, CH2), 1.83 (p, CH2). ^13^C NMR (101 MHz, DMSO-d6) δ 169.32, 165.30, 153.44, 45.00, 42.43, 37.14, 32.61, 23.31, 21.70, 20.32. The percent of platinum measured by ICP-MS is 10% (w/w).

### AA Polymer

Adipic acid (21.8mg, 0.15mmol, Sigma Aldrich) in DMSO (1mL) was added to a 20mL vial followed by DIC (5uL, 3 equiv.), DMAP (54.68mg, 3 equiv.), and oxoplatin (50mg, 1 equiv.). Reaction followed the same procedure as described for DTB Polymer. AA Polymer yield was 93.4mg. ^1^H NMR (400 MHz, DMSO-d6): δ 5.49 (d, NH3), 4.38 (m, NH3), 2.23 (d, CH2), 1.48 (d, CH2). ^13^C NMR (101 MHz, DMSO-d6) δ 169.72, 156.78, 153.57, 63.07, 44.89, 42.37, 40.66, 34.04, 24.60, 23.30, 21.67, 20.33. The percent of platinum measured by ICP-MS is 14% (w/w).

### C_16_-Met

Hexadecylamine (about 200mg, 0.83mmol, Sigma Aldrich, Cat # H7408-500G) and dicyandiamide (about 250mg, 4 equiv., Sigma Aldrich, Cat # D76609-25G) were added to an RB flask. 2M hydrochloric acid (HCl, 8mL) was added to the mixture and refluxed at 100°C for 24 hours.^23^ After cooling, the solution was filtered, and the precipitate was washed with excess water and diethyl ether to remove unreacted dicyandiamide and hexadecylamine, respectively. The resulting product was collected in a clean 20mL vial and left to air dry before storing at 4°C. ^1^H NMR of (400 MHz, DMSO-d6): δ 8.29 (s, 4H, NH), 2.97 (s, 2H, CH2), 1.76 (t, 2H, NH2), 1.25 (m, 28H, CH2), 0.87 (m, 3H, CH3). ^13^C NMR (101 MHz, DMSO-d6) δ 155.57, 154.45, 38.71, 31.29, 28.93, 28.84, 28.70, 28.54, 26.94, 25.83, 22.09, 13.96. ESI-MS calculated for m/z 325.32, found 326.3284 [M + H].

### C_6_-Met

Hexylamine (789μL, 5.9mmol, Sigma Aldrich, Cat # 8.04326.0100) and dicyandiamide (1.996g, 4 equiv.) were combined in a 100mL RB flask with 2M HCl (16mL) and refluxed at 100°C for 24 hours.^23^ After cooling, the solution was lyophilized. The solid product was washed with excess diethyl ether and the final product was confirmed with Thin layer chromatography (TLC, 1:10 MeOH: CHCl3). Product was stored at 4°C. ^1^H NMR (400 MHz, DMSO-d6): δ 10.18 (s, 4H, NH), 7.86 (s, 2H, NH2), 2.74 (s, 2H, CH2), 1.52 (q, 2H, CH2), 1.28 (d, 6H, CH2), 0.86 (m, 3H, CH3). ^13^C NMR (101 MHz, DMSO-d6) δ 155.66, 154.50, 38.72, 30.75, 26.89, 25.55, 21.92, 13.87.

### NP synthesis and characterization

Polymeric NPs were synthesized using previously described self-assembly nanoprecipitation methods.^26, 28^ Briefly, PLGA-PEG (50kDa – 5kDa MW, 75:25 L:G ratio, methyl terminated, PolySciTech, Cat # AK148) was dissolved in acetonitrile (ACN) at a concentration of 10mg/mL. 1mM solutions of DTB and AA Polymers were prepared in DMF. 30mg/mL solution of olaparib and 8mg/mL solutions of C_16_-Met and C_6_-Met in DMSO were all prepared. Single, dual, and triple drug NPs were synthesized using these drug solutions in different combinations. PLGA-PEG, cisplatin drug-polymer, olaparib, and metformin derivative were combined, maintaining a (v/v) ratio of 3:1 polymer:total drug content. The ratio between cisplatin polymer, olaparib, and metformin derivative was 1:2.5:2.5 (v/v/v). The polymer and drug solution was added dropwise to water (1:10 (v/v)), and spun (2h, RT) to allow self-assembly of NPs. Solutions were then transferred to 50mL centrifugal filter conical tubes (100kDa MWCO, VWR, Cat # MAP100C38) and centrifuged at roughly 3000rpm for 10 minutes. Once concentrated NPs were 1-10% of the starting volume (including the aqueous phase), they were collected in fresh microcentrifuge tubes (VWR) and stored at 4°C for further characterization and in vitro testing. Empty NPs were synthesized using the same outlined method, except drug volumes were replaced by solvent (DMF and/or DMSO) alone.

NP morphology was determined using TEM.^28^ A single droplet of concentrated NPs was added to carbon-coated copper grids (400 mesh, Ted Pella, Cat # 01822), the excess solution was wicked off onto filter paper, and stained with uranyl acetate (Electron Microscopy Sciences, Cat # 22409). Grids were washed with water and dryed (24h) and imaged using JEM-1400Flash TEM (120kv) system. NP size, polydispersity index (PDI) and surface charge (zeta potential) were all measured directly upon synthesis completion by dynamic light scattering using a ZetaSizer Nano Series machine (Malvern Panalytical, United Kingdom). To prepare samples, 10uL of concentrated NPs were diluted in 1mL sterile water. After 60 days of storage at 4°C, size and PDI were measured again to determine storage stability. As a surrogate for blood serum stability, NP size and PDI were measured following incubation with either 0, 5, or 10% FBS in sterile water at both 25°C and 37°C.

### Cell culture

Human epithelial SKOV-3 cells (ATCC, Manassas, Virginia, United States) were cultured in McCoy’s 5A medium (Cedarlane, Cat # 30-2007) supplemented with 10% heat inactivated fetal bovine serum (FBS, Corning), and 1% penicillin/streptomycin/L-Glutamine (Corning). Cells were grown in humidified incubators at 37°C with 5% CO_2_.

### MTT assays to determine drug synergy

Synergistic drug ratios were determined by MTT assays. SKOV-3 cells were seeded in 96-well plates and incubated with various concentrations of cisplatin, olaparib, metformin and all possible combinations for 72 hours. Drug media was replaced with MTT reagent (3-(4,5-Dimethylthiazol-2-yl)-2,5-Diphenyltetrazolium Bromide, Alfa Aesar, Cat # L11939) (5mg/mL in 1x phosphate buffered saline, PBS) in phenol red free culture media (Wisent); at a 1:9 (v/v) ratio. The plate was incubated for 30 minutes before lysing with 100uL DMSO. Purple formazan crystals were detected by optical density using a microplate reader at 570nm. Background absorbance was also detected at 630nm.

Percent cytotoxicity values were inputted into either CompuSyn software or Synergy Finder, two independent synergy calculating software. CompuSyn software generates CI values by using the Chou Talalay method to calculate synergy. CI < 1 is indicative of a synergistic concentration ratio between two or more agents in terms of cytotoxicity, whereas CI > 1 indicates an antagonistic relationship at specific concentration ratios^24^. Synergy Finder uses the highest single agent (HSA) reference method, in which synergy is denoted when the expected combination effect is equal or greater than the higher effect of individual drugs at the same concentrations^23^.

### Flow cytometry H2AX assays

SKOV-3 cells were seeded in 6-well plates (2×10^5^ cells/well) and treated with free cisplatin (positive control), DTB-NP, AA-NP, DTB-Ola-C_16_-Met-NP, E-NP (Empty-NP), or left untreated (negative control) for 24 hours. Cells were collected and centrifuged. The supernatant was discarded, and the cell pellet was resuspended and washed with 1x PBS and centrifuged again at 300g for 6 minutes. Supernatant was discarded again, and cell pellet was dislodged by light vortexing while adding ice cold 70% ethanol dropwise to fix and permeabilize the cells. Cells were incubated at −20°C for 60 minutes, then washed with cell staining buffer (BioLegend, Cat # 420201) 3 times before resuspending at a final concentration of roughly 10^6^ cells/100μL. Cells were stained with 5μL of FITC anti-H2A.X Phospho (Ser139) antibody (25μg/mL, BioLegend, Cat # 613403) for 1 hour covered from light at room temperature. Unbound antibody was removed by 2 more washes with cell staining buffer. Cell suspensions were then analyzed by the LSR Fortessa (BD Biosciences, California USA). The median fluorescence intensity (MFI) of the second peak from 2 independent replicates was analyzed and plotted as a bar graph.

### Protein extraction and western blot

For protein extraction, cells were washed with cold 1x PBS and collected into 1.5mL microcentrifuge tubes using cell scrapers (Fisher Scientific). Samples were centrifuged and cells were resuspended in 50μL RIPA lysis buffer (1M NaF, 1M β-glycerophosphate, 1M DTT, 1mg/mL leupeptin, 1mg/mL aprotinin, 0.2M benzamide, 150mM NaCl, 50mM Tris, 0.1% SDS, 1% Triton X-100, 0.5% Sodium Deoxycholate, pH 8.0, 1% pepstatin (1mg/mL), 1% PMSF (0.1M), and 0.625% sodium orthovanadate (100mM)), vortexed, and placed at −80°C until ready for western blotting. Before blotting, lysates were thawed and centrifuged at max speed for 10 minutes at 4°C to pellet cell debris. Protein concentration was quantified using a Bradford Lowry Reagent (Bio-Rad, Mississauga, Ontario, Canada). 20ug of protein lysate were loaded and separated on a 10% polyacrylamide gel and subsequently transferred to a polyvinylidene difluoride (PVDF) membrane (Bio-Rad). Membranes were blocked in either 5% bovine serum albumin (BSA, Sigma Aldrich) or skim milk in 1x tris-buffered saline with 0.1% Tween-20 (TBST) for 1 hour at room temperature. Membranes were washed with TBST before incubating with primary antibody overnight at 4°C. Three washes were performed again before secondary antibody incubation for 1 hour at room temperature. Membranes were washed 3 more times before imaging with a 1:1 (v/v) ratio of Clarity Western enhanced chemiluminescence and peroxide solutions (Bio-Rad, Cat # 1705061). Primary and secondary antibodies used include: phospho-AMPKα (Thr172) (1:1000 in BSA, Cell Signaling Technology, Cat # 2535S), AMPKα (1:1000 in BSA, Cell Signaling Technology, Cat # 5831S), PARP (1:1000 in milk, Cell Signaling Technology, Cat # 9542S), cleaved PARP (Asp214) (1:1000 in milk, Cell Signaling Technology, Cat # 9541S), Caspase-3 (1:1000 in milk, Cell Signaling Technology, Cat # 9662S), cleaved Caspase-3 (Asp175) (1:1000, Cell Signaling Technology, Cat # 9664T), GAPDH (1:1000, Thermo Fisher, Cat # MA5-15738), anti-rabbit IgG, HRP-linked (1:2000, Cell Signaling Technology, Cat # 7074S), and anti-mouse IgG, HRP-linked (1:2000, Cell Signaling Technology, Cat # 7076S).

### Cell viability

Cell viability following NP treatment was assessed using MTT assays as previously described.

### Statistical analysis

Data is presented as means ± standard error of the mean (SEM). Three technical replicates were performed for every biological replicate. Where appropriate, statistical analysis (paired Student’s t test) was performed using GraphPad prism. Statistical significance was defined as p < 0.05.

## Author contributions

All authors were involved in data collection, analysis, and final editing.

## Supporting information

SI with raw data

## Acknowledgment

The authors would like to thank Vera Tang from the uOttawa Flow Cytometry and Virometry core for technical support.

## Supporting Information

The Supporting Information contains, ^1^H, ^13^C NMRs, cell viability and synergy data of ID8;p53^-/-^;BRCA1^-/-^, A2780-S and A2780-CP cells, size and PDI of dual drug NPs, and additional flow cytometry and western blot data with cisplatin prodrugs and metformin derivatives in SKOV-3 cells.

