## Supplementary material for "Development of 3-in-1 nanotherapeutic strategies for ovarian cancer": SI

*Corresponding author

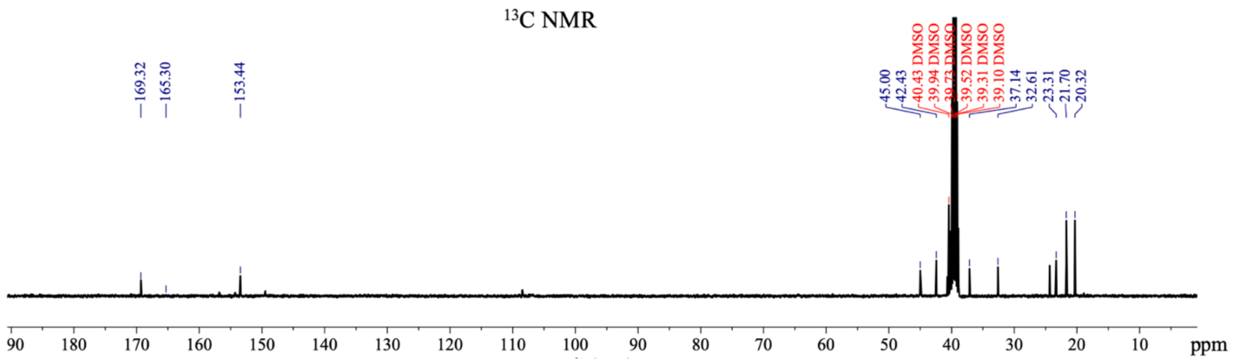


Figure S1. $^{13}C$ NMR spectrum of DTB Polymer in DMSO-d6.

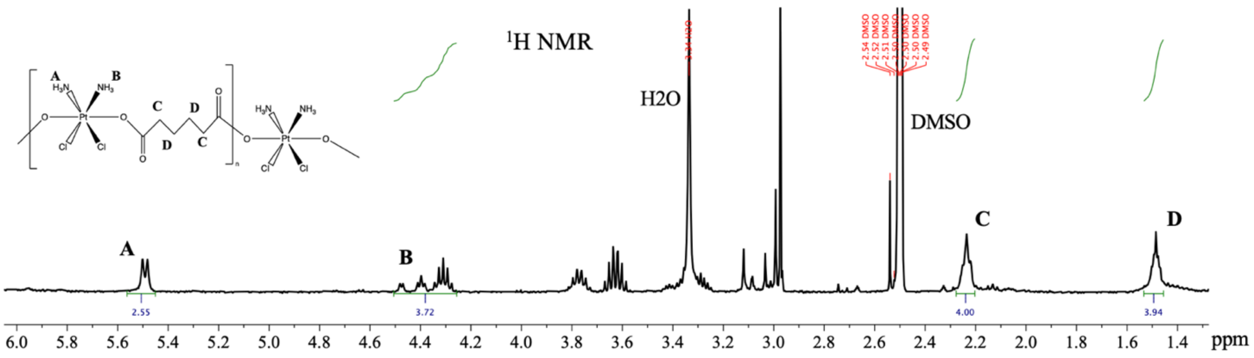


Figure S2. $^{1}H$ NMR spectrum of AA Polymer in DMSO-d6.

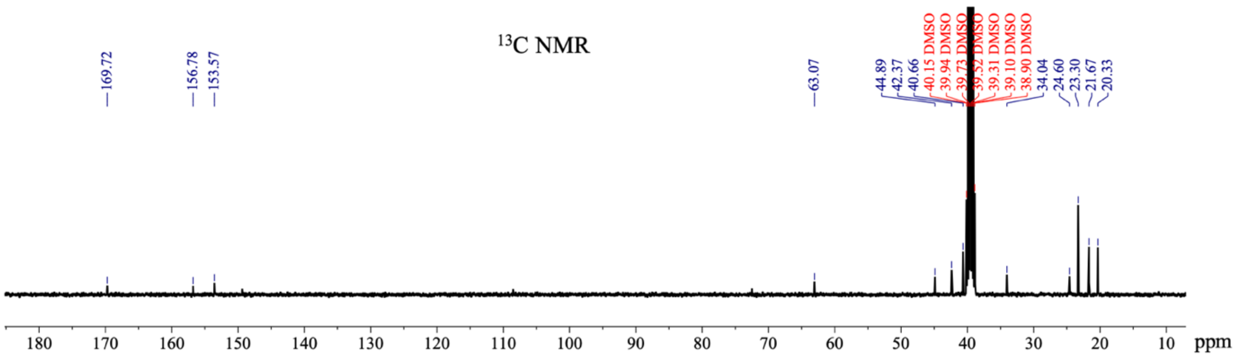


Figure S3. $^{13}C$ NMR spectrum of AA Polymer in DMSO-d6.

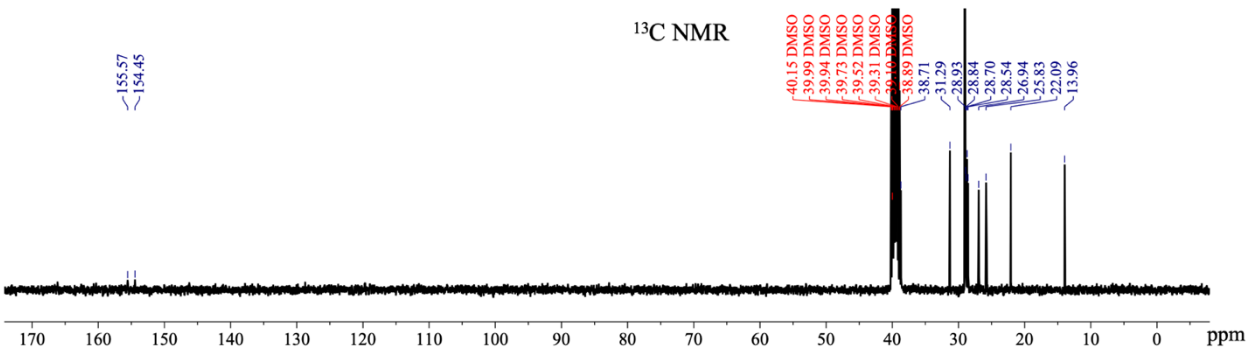


Figure S4. $^{13}C$ NMR spectrum of $C_{16}$-Met in DMSO-d6.

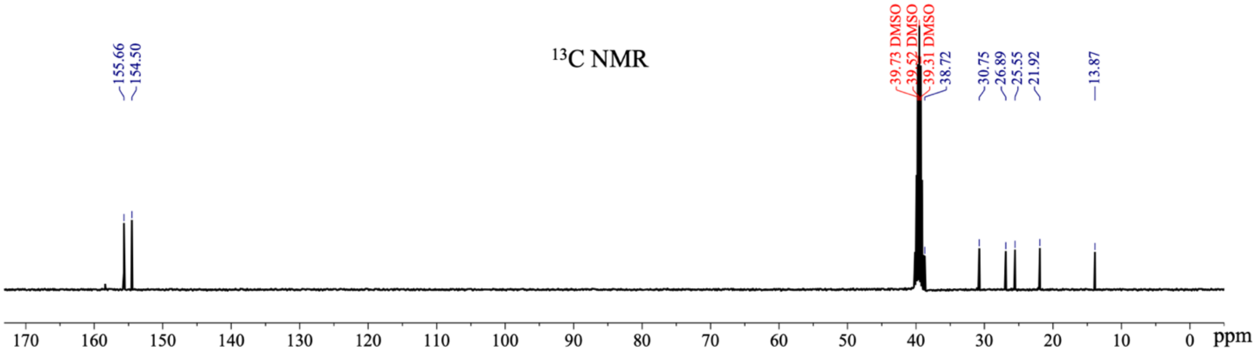


Figure S5. $^{13}C$ NMR spectrum of $C_6$-Met in DMSO-d6.

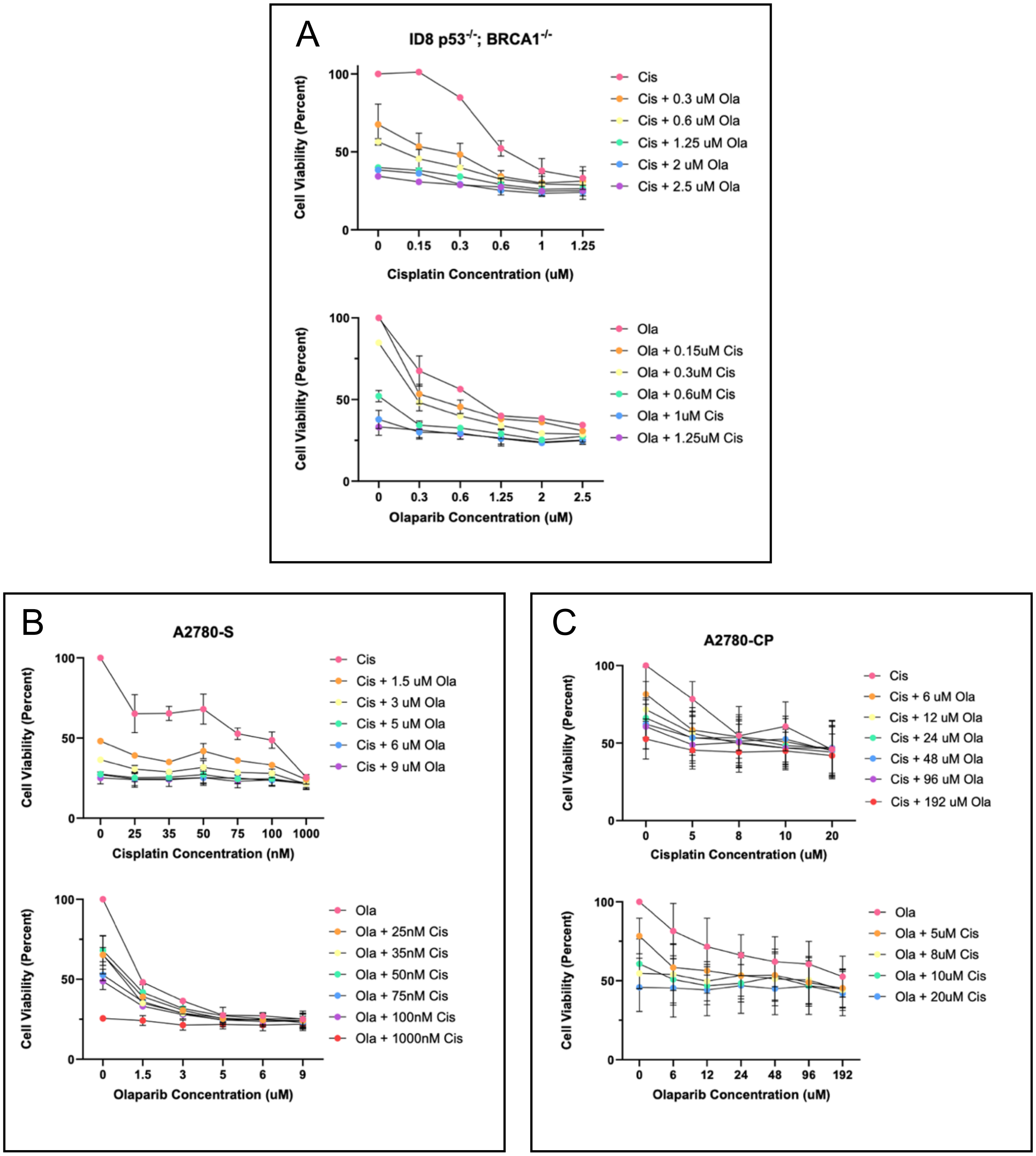


Figure S6. Cell viability following single agent and combination treatment with cisplatin and olaparib. (A) ID8 $p53^{-/-}$; $BRCA1^{-/-}$, (B) A2780-S, and (C) A2780-CP cell viability following treatment with cisplatin, olaparib and the two agents in combination for 72 hours plotted either against cisplatin concentration (top) or olaparib concentration (bottom). Data measured by MTT assays. Data presented as means ± SEM (n=2).

A

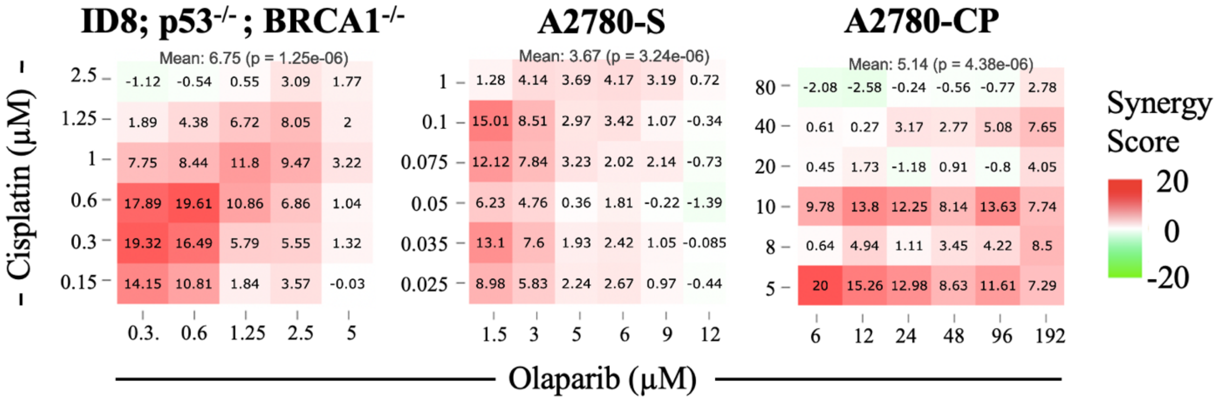


B

| | ID8; p53-/- ; BRCA1-/- | A2780-S | A2780-CP |
|---|---|---|---|
| Cisplatin : Olaparib (µM) | 0.15-1.25 : 0.3-2.5 | 0.025-0.1 : 1.5-6 | 5-40 : 6-192 |
| Combination Index | 0.59 – 0.96 | 0.52 – 0.96 | 0.38 – 0.98 |

Figure S7. Synergistic ratios between cisplatin and olaparib across different ovarian cancer cell lines. MTT data was inputted into either (A) Synergy Finder or (B) CompuSyn to obtain synergy scores and combination index (CI) values, respectively for ID8 p53$^{-/-}$; BRCA1$^{-/-}$ A2780-S and A2780-CP cells (n=2). The synergy scores were calculated using the HSA reference model. CompuSyn calculates CI values using the Chou Talalay method.

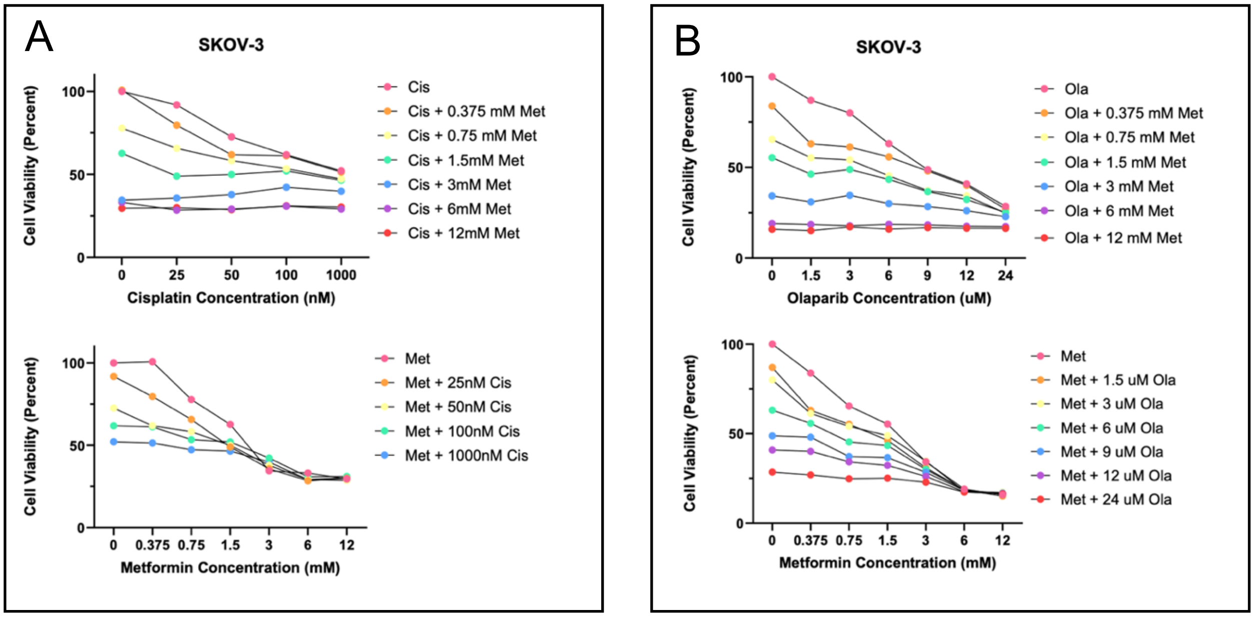


Figure S8. Cell viability following single agent and combination treatment with either cisplatin and olaparib or olaparib and metformin. (A) Cell viability following treatment with cisplatin, metformin and the two agents in combination for 72 hours plotted either against cisplatin concentration (top) or metformin concentration (bottom) (n=1). (B) Cell viability following treatment with olaparib, metformin and the two agents in combination for 72 hours plotted either against olaparib concentration (top) or metformin concentration (bottom) (n=1). Data measured by MTT assays.

| | **ID8; p53$^{-/-}$; BRCA1$^{-/-}$** | **A2780-S** | **A2780-CP** |
|---|---|---|---|
| Cisplatin : Olaparib : Metformin (μM) | 0.156-0.625 : 0.3-1.25 : 3000-6000 | 0.025-0.1 : 1.5-6 : 250-1000 | 5-20 : 6-24 : 3000-6000 |
| Combination Index | 0.73 – 0.96 | 0.50 – 0.94 | 0.49 – 0.96 |

Figure S9. Synergistic ratio between cisplatin, olaparib, and metformin across different ovarian cancer cell lines. MTT data was inputted into CompuSyn to obtain combination index (CI) values for ID8 p53$^{-/-}$; BRCA1$^{-/-}$, A2780-S, and A2780-CP cells. CI<1 indicates synergy, whereas CI>1 represents antagonism (n=1).

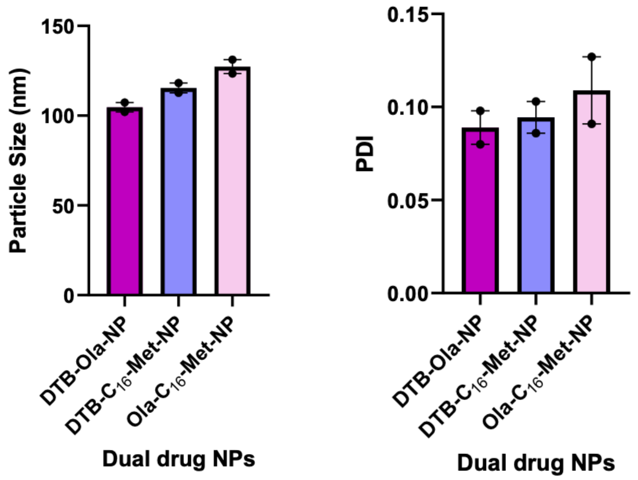


Figure S10. Size and PDI of dual drug NPs measured by dynamic light scattering (n=2 or 1)

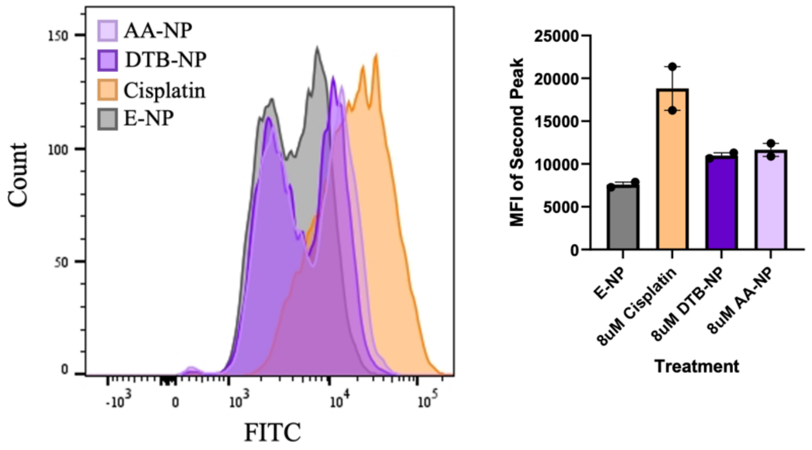


Figure S11. Cisplatin polymers induce DNA damage in SKOV-3 cells. Representative flow cytometry histogram of FITC-tagged γH2AX induced by cisplatin in SKOV-3 cells (left) and MFI (right) following E-NP, 8μM cisplatin, 8μM DTB-NP and 8μM AA-NP treatment for 24 hours. Data is presented as means ± SEM (n=2).

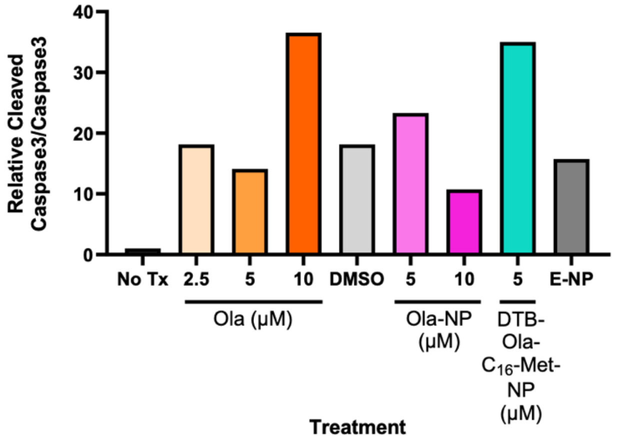


Figure S12. Quantification of relative cleaved caspase 3/caspase 3 ratio from from figure 6B. Data normalized to No Tx control, which is set to 1. Data analyzed by Fiji (n=1).

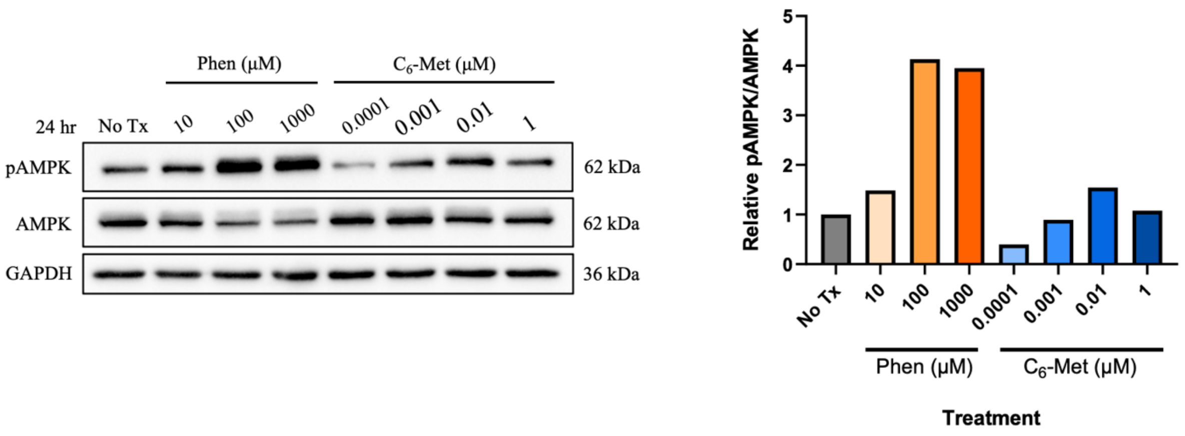


Figure S13. $C_6$-Met derivative activates AMPK in SKOV-3 cells. Western blot of phosphorylated AMPK (pAMPK) in SKOV-3 cells after $C_6$-Met (left) treatment for 24 hours. Phenformin was used as a positive control. Relative pAMPK/AMPK ratio (right) quantified using Fiji. Data normalized to No Tx control, which is set to 1 (n=1).
