## Supplementary material for "Development of 3-in-1 nanotherapeutic strategies for ovarian cancer": SI with raw data

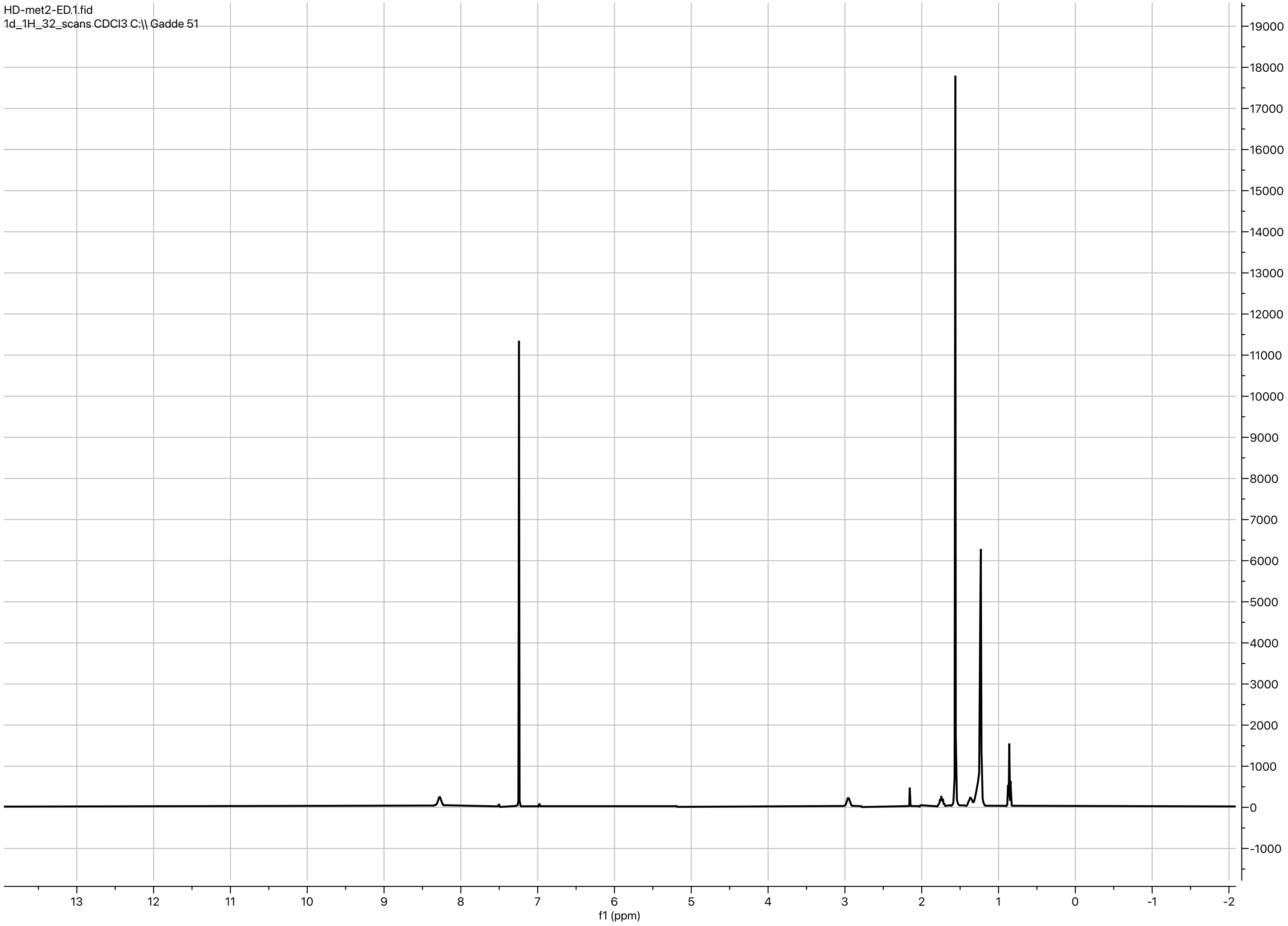


Fig. 2 from MS

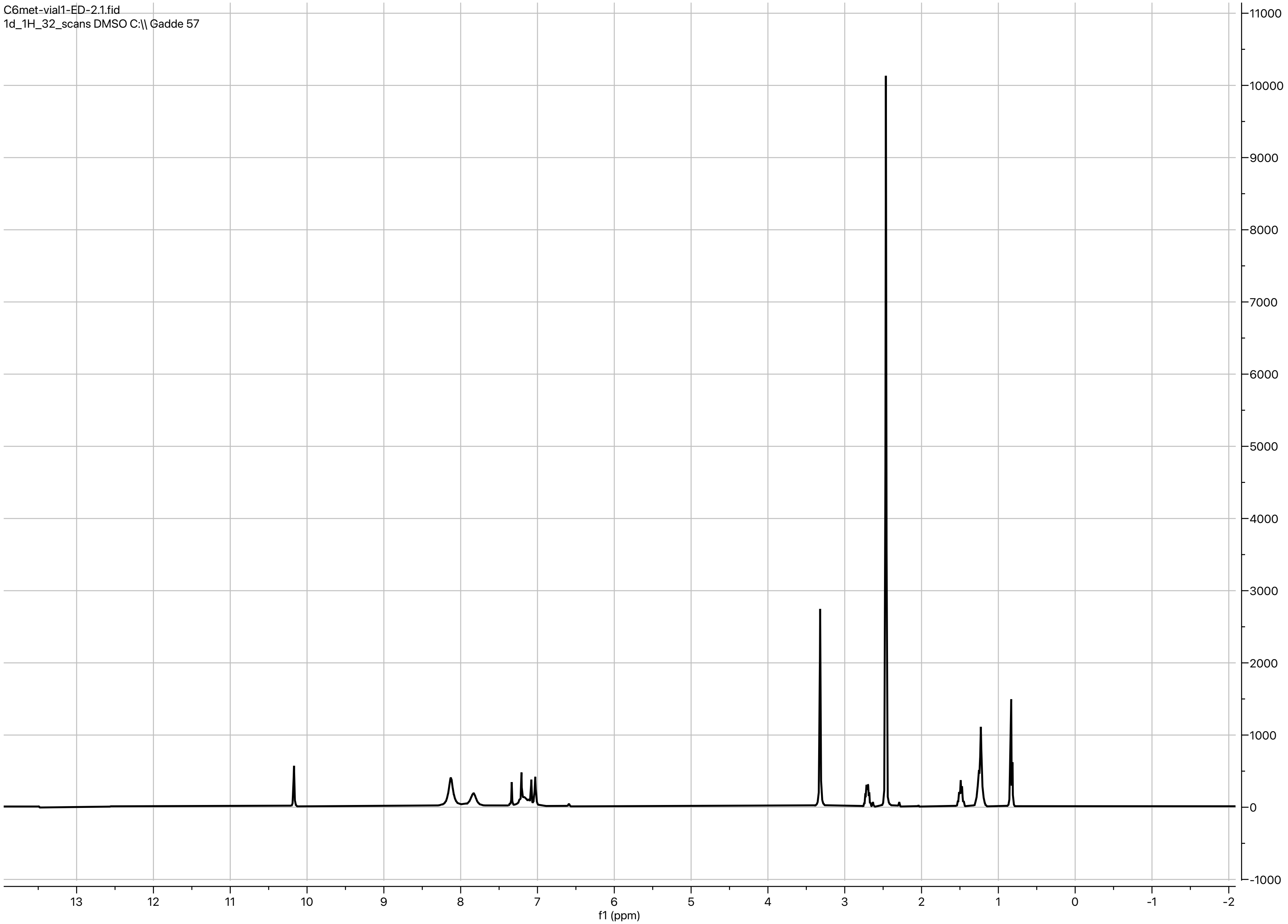


Fig. 2 from MS

### Synergy Finder Data from MTT Assays

| Drug1: | Olaparib | | | | | |
|---|---|---|---|---|---|---|
| Drug2: | Cisplatin | | | | | | |
| ConcUnit: | uM | | | | | | |
| | 0 | 0.156 | 0.3 | 0.6 | 1 | 1.25 | 2.5 |
| 12 | 42.87 | 43.05 | 46.8 | 46.44 | 48.94 | 49.48 | 47.51 |
| 9 | 39.3 | 31.8 | 44.48 | 45.01 | 45.55 | 46.09 | 49.3 |
| 6 | 33.05 | 26.45 | 34.3 | 42.69 | 40.73 | 45.19 | 47.16 |
| 5 | 25.55 | 32.52 | 38.94 | 40.73 | 36.98 | 40.55 | 46.09 |
| 3 | 13.77 | 31.62 | 36.44 | 48.58 | 39.12 | 43.23 | 54.3 |
| 1.5 | 7.88 | 23.06 | 29.13 | 42.87 | 37.87 | 51.26 | 47.33 |
| 0 | 0 | 5.02 | 11.09 | 40.37 | 33.23 | 46.62 | 53.76 |

| | | | | | | | |
|---|---|---|---|---|---|---|---|
| Drug1: | Metformin | | | | | | |
| Drug2: | Cisplatin | | | | | | |
| ConcUnit: | uM | | | | | | |
| | 0 | 0.025 | 0.035 | 0.05 | 0.075 | 0.1 | 1 |
| 12000 | 70.43 | 70.09 | 71.11 | 71.22 | 71.78 | 68.96 | 69.75 |
| 6000 | 66.82 | 71.56 | 68.96 | 70.88 | 70.2 | 69.19 | 70.88 |
| 3000 | 65.46 | 64.33 | 62.64 | 62.19 | 65.24 | 57.79 | 60.16 |
| 1500 | 37.36 | 51.13 | 55.19 | 50.11 | 56.09 | 47.97 | 53.61 |
| 750 | 22.23 | 34.31 | 45.49 | 41.76 | 48.53 | 46.61 | 52.71 |
| 375 | -0.68 | 20.43 | 48.76 | 38.15 | 41.08 | 38.83 | 48.65 |
| 0 | 0 | 8.24 | 45.94 | 27.43 | 46.73 | 38.15 | 47.86 |

| | | | | | | | |
|---|---|---|---|---|---|---|---|
| Drug1: | Metformin | | | | | | |
| Drug2: | Olaparib | | | | | | |
| ConcUnit: | uM | | | | | | |
| | 0 | 1.5 | 3 | 6 | 9 | 12 | 24 |
| 12000 | 84.11 | 84.9 | 82.76 | 84.03 | 83.24 | 83.55 | 83.63 |
| 6000 | 80.93 | 81.49 | 82.2 | 81.41 | 81.73 | 82.44 | 82.6 |
| 3000 | 65.68 | 69.01 | 65.36 | 69.97 | 71.64 | 73.86 | 77.12 |
| 1500 | 44.62 | 53.68 | 51.14 | 56.62 | 63.37 | 67.74 | 74.89 |
| 750 | 34.53 | 44.7 | 45.9 | 54.63 | 62.82 | 65.68 | 75.21 |
| 375 | 16.1 | 37 | 38.74 | 44.23 | 51.93 | 59.88 | 73.07 |
| 0 | 0 | 12.92 | 19.99 | 36.84 | 51.22 | 59.16 | 71.48 |

Fig. 3 from MS

### CompuSyn Data

#### CI Data for Non-Constant Combo: C+O (Cis+Ola)

| Dose Cis | Dose Ola | Effect | CI |
|---|---|---|---|
| 2.5 | 12.0 | 0.4751 | 2.64049 |
| 2.5 | 9.0 | 0.493 | 2.25439 |
| 2.5 | 6.0 | 0.4716 | 2.18595 |
| 2.5 | 5.0 | 0.4609 | 2.18490 |
| 2.5 | 3.0 | 0.543 | 1.51566 |
| 2.5 | 1.5 | 0.4733 | 1.80985 |
| 1.25 | 12.0 | 0.4948 | 1.68042 |
| 1.25 | 9.0 | 0.4609 | 1.64098 |
| 1.25 | 6.0 | 0.4519 | 1.43258 |
| 1.25 | 5.0 | 0.4055 | 1.58779 |
| 1.25 | 3.0 | 0.4323 | 1.25489 |
| 1.25 | 1.5 | 0.5126 | 0.84238 |
| 1.0 | 12.0 | 0.4894 | 1.55297 |
| 1.0 | 9.0 | 0.4555 | 1.49300 |
| 1.0 | 6.0 | 0.4073 | 1.46728 |
| 1.0 | 5.0 | 0.3698 | 1.56698 |
| 1.0 | 3.0 | 0.3912 | 1.22919 |
| 1.0 | 1.5 | 0.3787 | 1.11563 |
| 0.625 | 12.0 | 0.4644 | 1.43539 |
| 0.625 | 9.0 | 0.4501 | 1.24736 |
| 0.625 | 6.0 | 0.4269 | 1.06930 |
| 0.625 | 5.0 | 0.4073 | 1.04536 |
| 0.625 | 3.0 | 0.4858 | 0.63582 |
| 0.625 | 1.5 | 0.4287 | 0.63552 |
| 0.3125 | 12.0 | 0.468 | 1.20209 |
| 0.3125 | 9.0 | 0.4448 | 1.03816 |
| 0.3125 | 6.0 | 0.343 | 1.12525 |
| 0.3125 | 5.0 | 0.3894 | 0.83257 |
| 0.3125 | 3.0 | 0.3644 | 0.67334 |
| 0.3125 | 1.5 | 0.2913 | 0.65860 |
| 0.15625 | 12.0 | 0.4305 | 1.25339 |
| 0.15625 | 9.0 | 0.318 | 1.49401 |
| 0.15625 | 6.0 | 0.2645 | 1.33338 |
| 0.15625 | 5.0 | 0.3252 | 0.88621 |
| 0.15625 | 3.0 | 0.3162 | 0.62652 |
| 0.15625 | 1.5 | 0.2306 | 0.59620 |

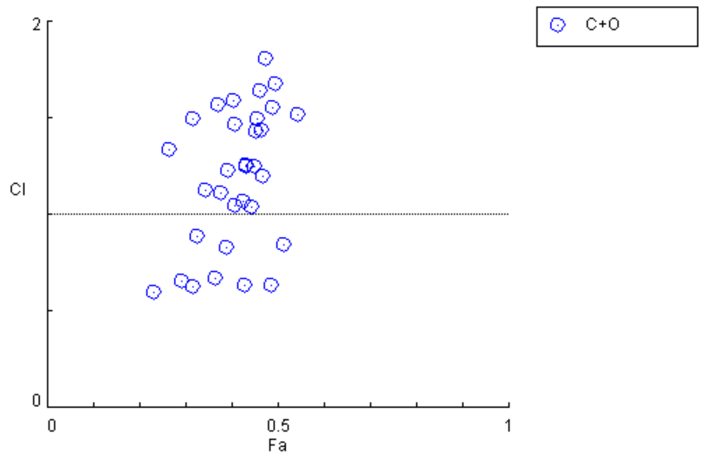


Fig. 3 from MS

### CompuSyn Data

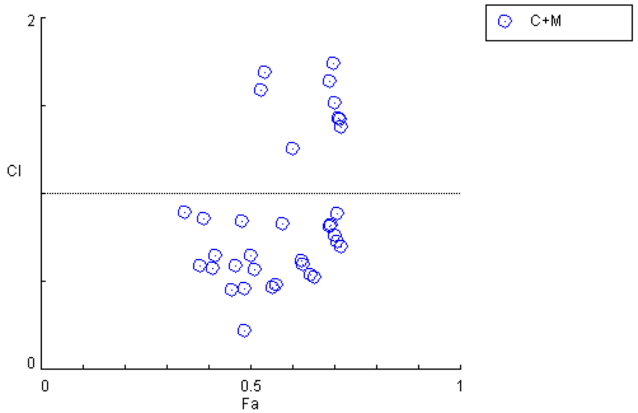


#### CI Data for Non-Constant Combo: C+M (Cis+Met)

| Dose Cis | Dose Met | Effect | CI |
|---|---|---|---|
| 1.0 | 12000.0 | 0.6975 | 1.73958 |
| 1.0 | 6000.0 | 0.7088 | 0.88855 |
| 1.0 | 3000.0 | 0.6016 | 1.26046 |
| 1.0 | 1500.0 | 0.5361 | 1.69062 |
| 1.0 | 750.0 | 0.5271 | 1.59372 |
| 1.0 | 375.0 | 0.4865 | 2.23724 |
| 0.1 | 12000.0 | 0.6896 | 1.64330 |
| 0.1 | 6000.0 | 0.6919 | 0.82065 |
| 0.1 | 3000.0 | 0.5779 | 0.82776 |
| 0.1 | 1500.0 | 0.4797 | 0.84418 |
| 0.1 | 750.0 | 0.4661 | 0.59199 |
| 0.1 | 375.0 | 0.3883 | 0.85966 |
| 0.075 | 12000.0 | 0.7178 | 1.37698 |
| 0.075 | 6000.0 | 0.702 | 0.76648 |
| 0.075 | 3000.0 | 0.6524 | 0.52743 |
| 0.075 | 1500.0 | 0.5609 | 0.47978 |
| 0.075 | 750.0 | 0.4853 | 0.45971 |
| 0.075 | 375.0 | 0.4108 | 0.57714 |
| 0.05 | 12000.0 | 0.7122 | 1.42237 |
| 0.05 | 6000.0 | 0.7088 | 0.73053 |
| 0.05 | 3000.0 | 0.6219 | 0.61786 |
| 0.05 | 1500.0 | 0.5011 | 0.64500 |
| 0.05 | 750.0 | 0.4176 | 0.64665 |
| 0.05 | 375.0 | 0.3815 | 0.58795 |
| 0.035 | 12000.0 | 0.7111 | 1.42973 |
| 0.035 | 6000.0 | 0.6896 | 0.81846 |
| 0.035 | 3000.0 | 0.6264 | 0.59559 |
| 0.035 | 1500.0 | 0.5519 | 0.46508 |
| 0.035 | 750.0 | 0.4549 | 0.45428 |
| 0.035 | 375.0 | 0.4876 | 0.22101 |
| 0.025 | 12000.0 | 0.7009 | 1.52041 |
| 0.025 | 6000.0 | 0.7156 | 0.69622 |
| 0.025 | 3000.0 | 0.6433 | 0.53778 |
| 0.025 | 1500.0 | 0.5113 | 0.56768 |
| 0.025 | 750.0 | 0.3431 | 0.89522 |
| 0.025 | 375.0 | 0.2043 | 2.51460 |

Fig. 3 from MS

CompuSyn Data

### CI Data for Non-Constant Combo: O+M (ola+met)

| Dose ola | Dose met | Effect | CI |
|---|---|---|---|
| 24.0 | 12000.0 | 0.8363 | 1.87978 |
| 24.0 | 6000.0 | 0.826 | 1.31061 |
| 24.0 | 3000.0 | 0.7712 | 1.32817 |
| 24.0 | 1500.0 | 0.7489 | 1.20824 |
| 24.0 | 750.0 | 0.7521 | 1.04812 |
| 24.0 | 375.0 | 0.7307 | 1.08539 |
| 12.0 | 12000.0 | 0.8355 | 1.60942 |
| 12.0 | 6000.0 | 0.8244 | 1.02233 |
| 12.0 | 3000.0 | 0.7386 | 1.08976 |
| 12.0 | 1500.0 | 0.6774 | 1.04833 |
| 12.0 | 750.0 | 0.6568 | 0.92119 |
| 12.0 | 375.0 | 0.5988 | 1.02167 |
| 9.0 | 12000.0 | 0.8324 | 1.57430 |
| 9.0 | 6000.0 | 0.8173 | 0.99354 |
| 9.0 | 3000.0 | 0.7164 | 1.08191 |
| 9.0 | 1500.0 | 0.6337 | 1.07290 |
| 9.0 | 750.0 | 0.6282 | 0.84181 |
| 9.0 | 375.0 | 0.5193 | 1.08944 |
| 6.0 | 12000.0 | 0.8403 | 1.41706 |
| 6.0 | 6000.0 | 0.8141 | 0.93423 |
| 6.0 | 3000.0 | 0.6997 | 1.02689 |
| 6.0 | 1500.0 | 0.5662 | 1.16306 |
| 6.0 | 750.0 | 0.5463 | 0.89688 |
| 6.0 | 375.0 | 0.4423 | 1.06542 |
| 3.0 | 12000.0 | 0.8276 | 1.48115 |
| 3.0 | 6000.0 | 0.822 | 0.80880 |
| 3.0 | 3000.0 | 0.6536 | 1.09091 |
| 3.0 | 1500.0 | 0.5114 | 1.14037 |
| 3.0 | 750.0 | 0.459 | 0.88813 |
| 3.0 | 375.0 | 0.3874 | 0.83597 |
| 1.5 | 12000.0 | 0.849 | 1.22890 |
| 1.5 | 6000.0 | 0.8149 | 0.80882 |
| 1.5 | 3000.0 | 0.6901 | 0.84758 |
| 1.5 | 1500.0 | 0.5368 | 0.89164 |
| 1.5 | 750.0 | 0.447 | 0.73794 |
| 1.5 | 375.0 | 0.37 | 0.63773 |

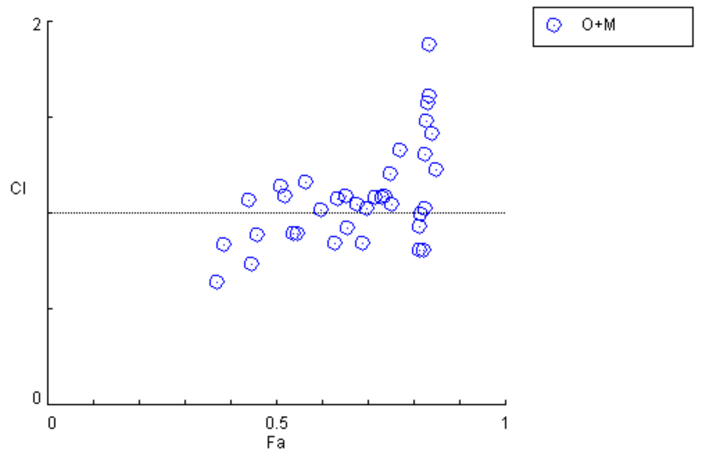


Fig. 3 from MS

### CompuSyn Data

#### CI Data for Non-Constant Combo: C+O+M (Cis+Ola+Met)

| Dose Cis | Dose Ola | Dose Met | Effect | CI |
|---|---|---|---|---|
| 0.2 | 12.0 | 200.0 | 0.5815 | 1.11178 |
| 0.2 | 5.0 | 200.0 | 0.3711 | 2.10958 |
| 0.2 | 1.5 | 200.0 | 0.3259 | 1.69021 |
| 0.05 | 12.0 | 200.0 | 0.58 | 1.03597 |
| 0.05 | 5.0 | 200.0 | 0.342 | 2.10562 |
| 0.05 | 1.5 | 200.0 | 0.215 | 2.45544 |
| 0.0125 | 12.0 | 200.0 | 0.5692 | 1.07654 |
| 0.0125 | 5.0 | 200.0 | 0.2884 | 2.79543 |
| 0.0125 | 1.5 | 200.0 | 0.2089 | 2.27314 |
| 0.2 | 12.0 | 100.0 | 0.5899 | 0.92645 |
| 0.2 | 5.0 | 100.0 | 0.3512 | 2.08885 |
| 0.2 | 1.5 | 100.0 | 0.2448 | 2.42857 |
| 0.05 | 12.0 | 100.0 | 0.5601 | 1.00824 |
| 0.05 | 5.0 | 100.0 | 0.2961 | 2.45438 |
| 0.05 | 1.5 | 100.0 | 0.1637 | 3.14583 |
| 0.0125 | 12.0 | 100.0 | 0.57 | 0.92888 |
| 0.0125 | 5.0 | 100.0 | 0.2609 | 2.97735 |
| 0.0125 | 1.5 | 100.0 | 0.0926 | 6.64820 |
| 0.2 | 12.0 | 50.0 | 0.583 | 0.89665 |
| 0.2 | 5.0 | 50.0 | 0.4009 | 1.41645 |
| 0.2 | 1.5 | 50.0 | 0.3856 | 0.83597 |
| 0.05 | 12.0 | 50.0 | 0.5937 | 0.76154 |
| 0.05 | 5.0 | 50.0 | 0.2992 | 2.23158 |
| 0.05 | 1.5 | 50.0 | 0.2081 | 1.84166 |
| 0.0125 | 12.0 | 50.0 | 0.5968 | 0.72809 |
| 0.0125 | 5.0 | 50.0 | 0.3015 | 2.05955 |
| 0.0125 | 1.5 | 50.0 | 0.2471 | 1.12538 |

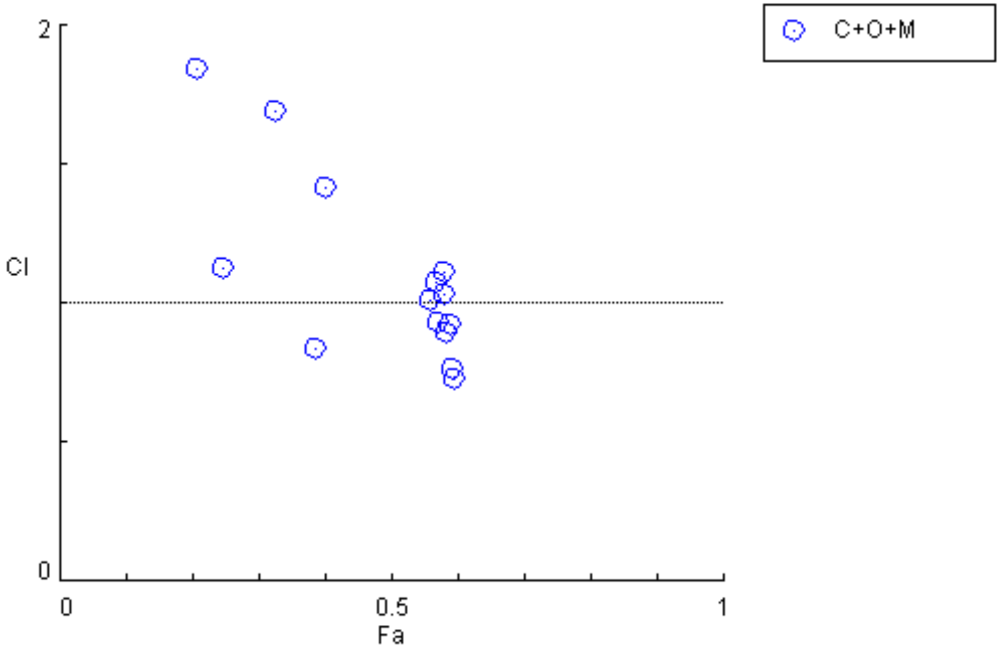


Fig. 3 from MS

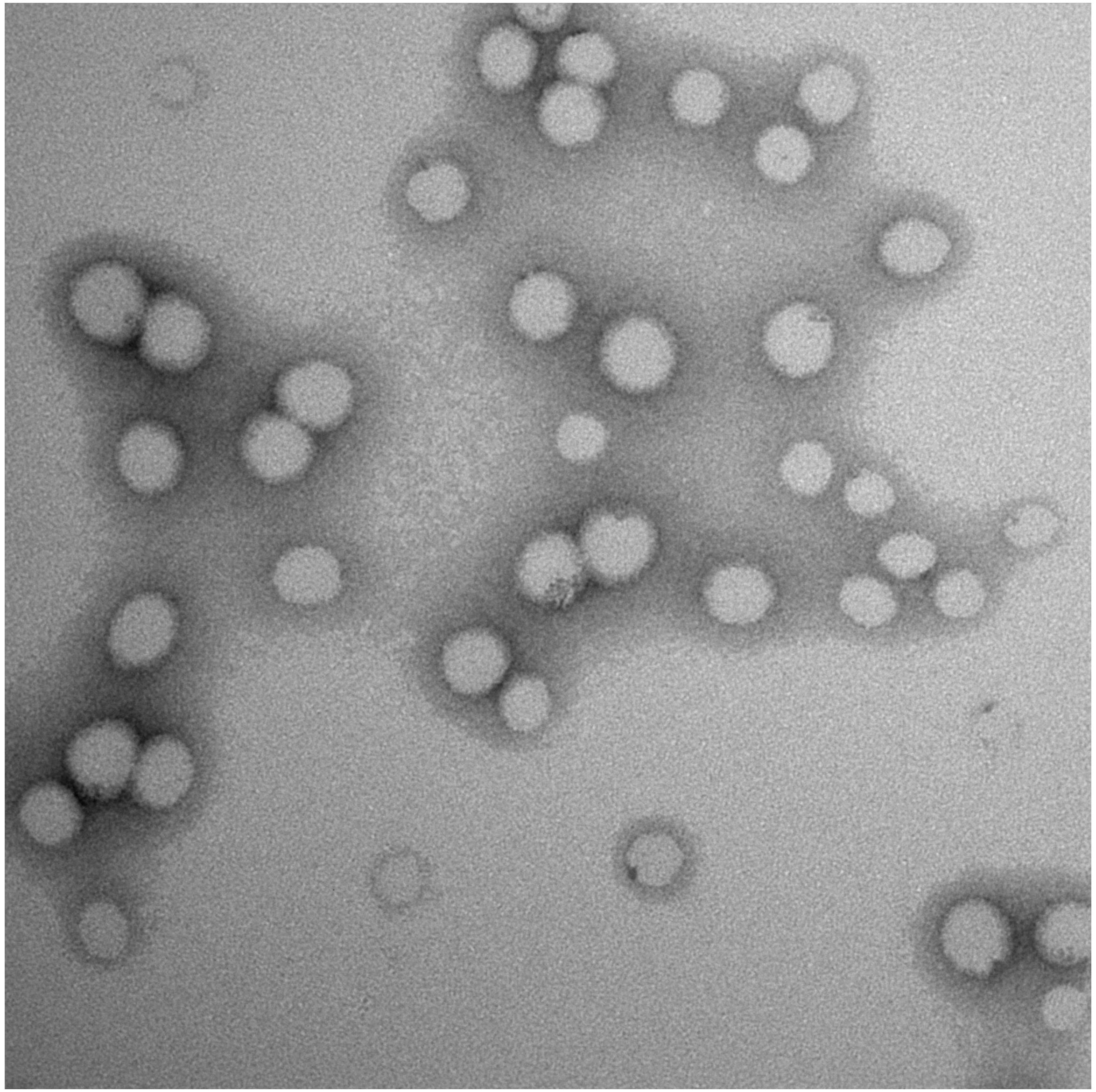

Fig. 4 from MS

### Ungated Flow Cytometry Data (first repeat)

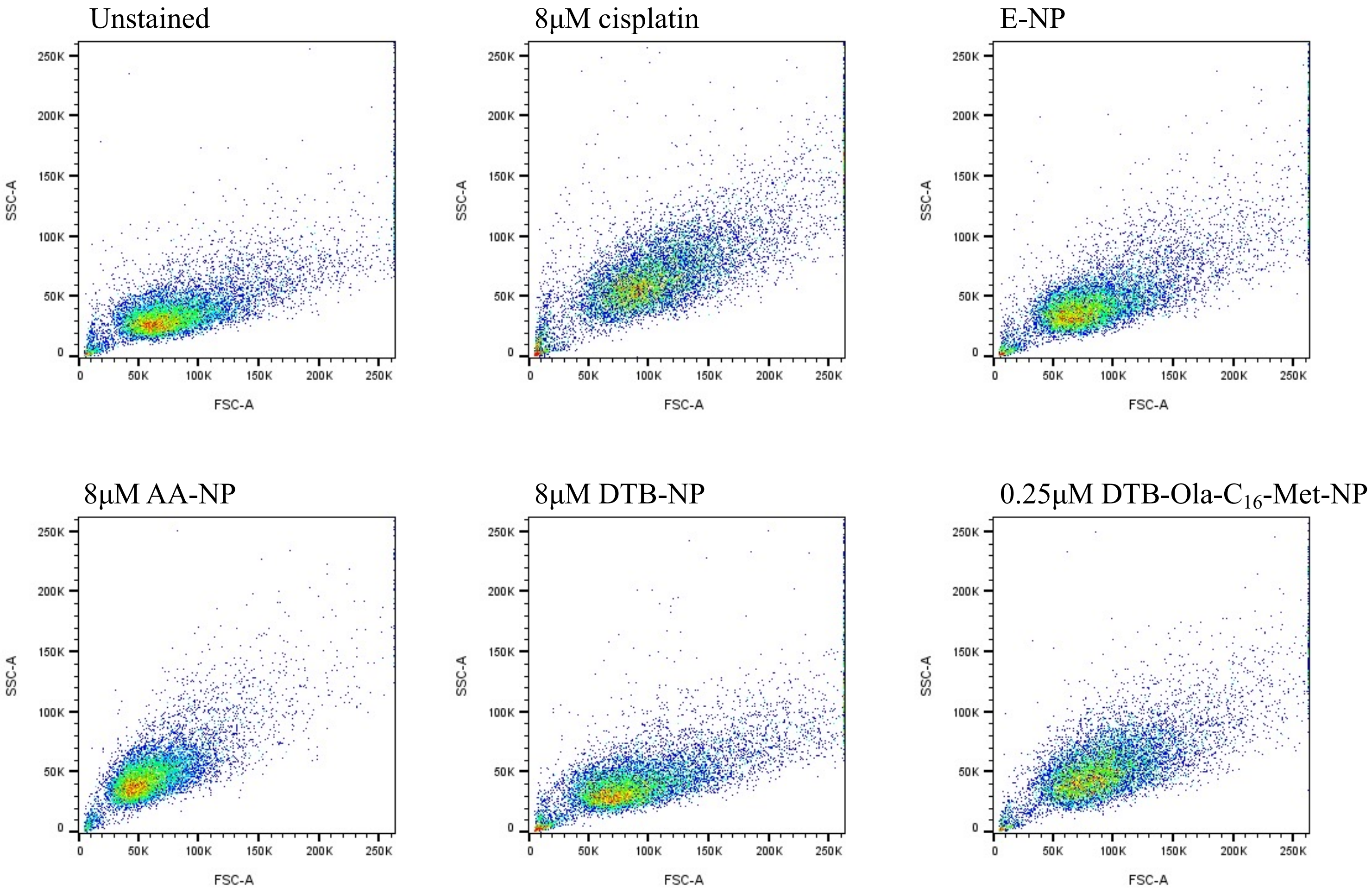


Fig. 6 from MS

### Gated Flow Cytometry Data (first repeat)

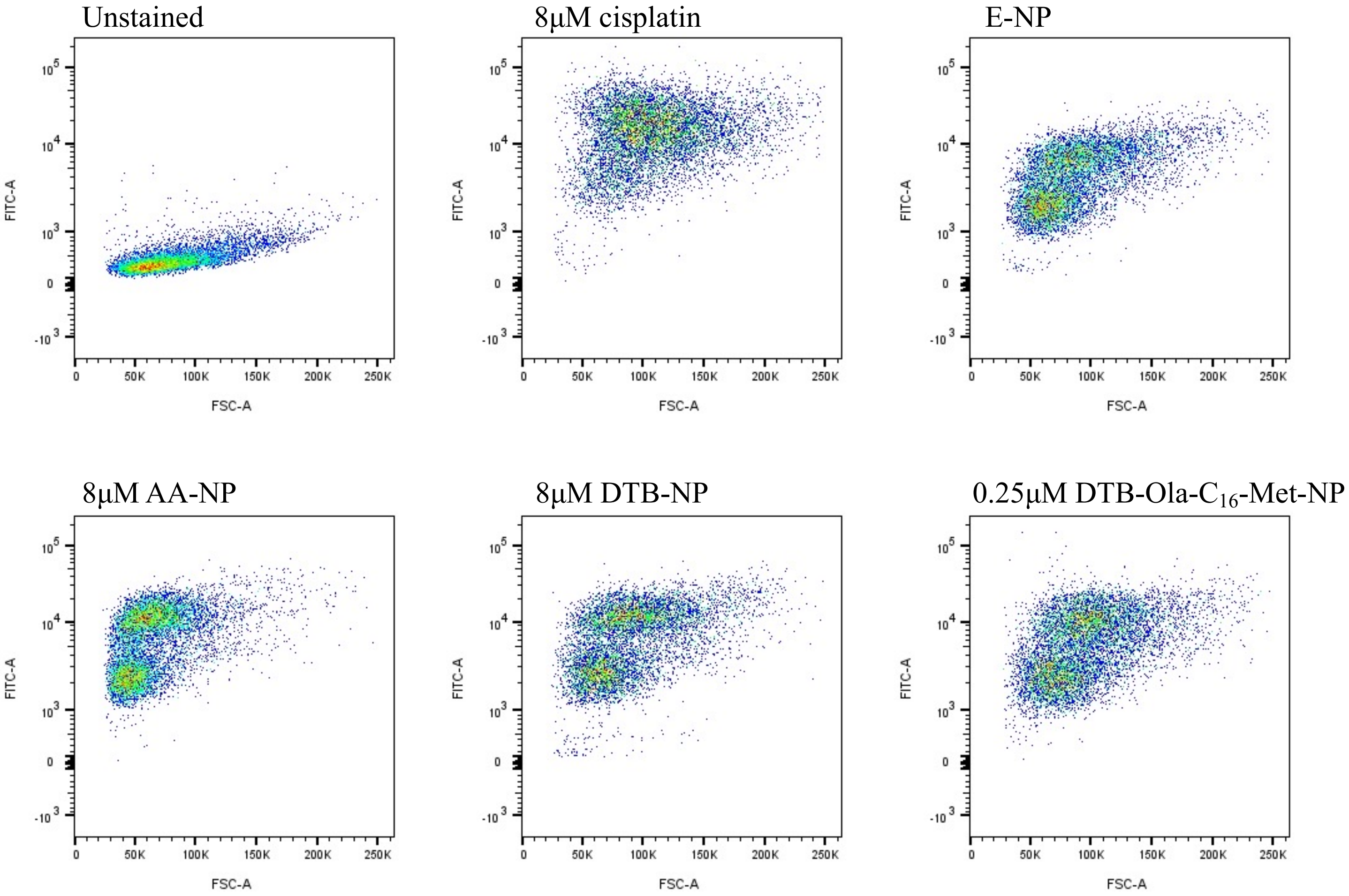


Fig. 6 from MS

### Ungated Flow Cytometry Data (second repeat)

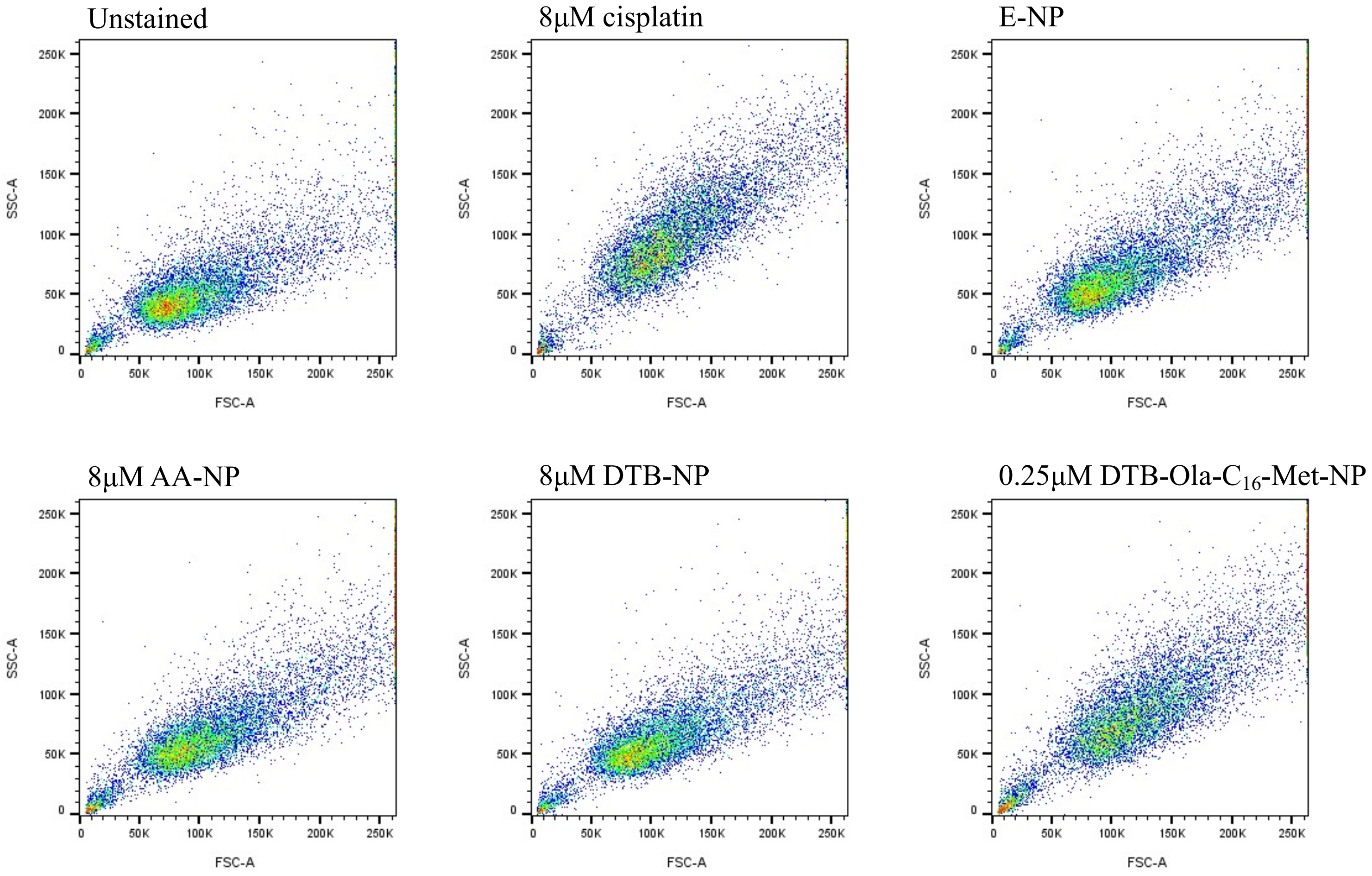


Fig. 6 from MS

### Gated Flow Cytometry Data (second repeat)

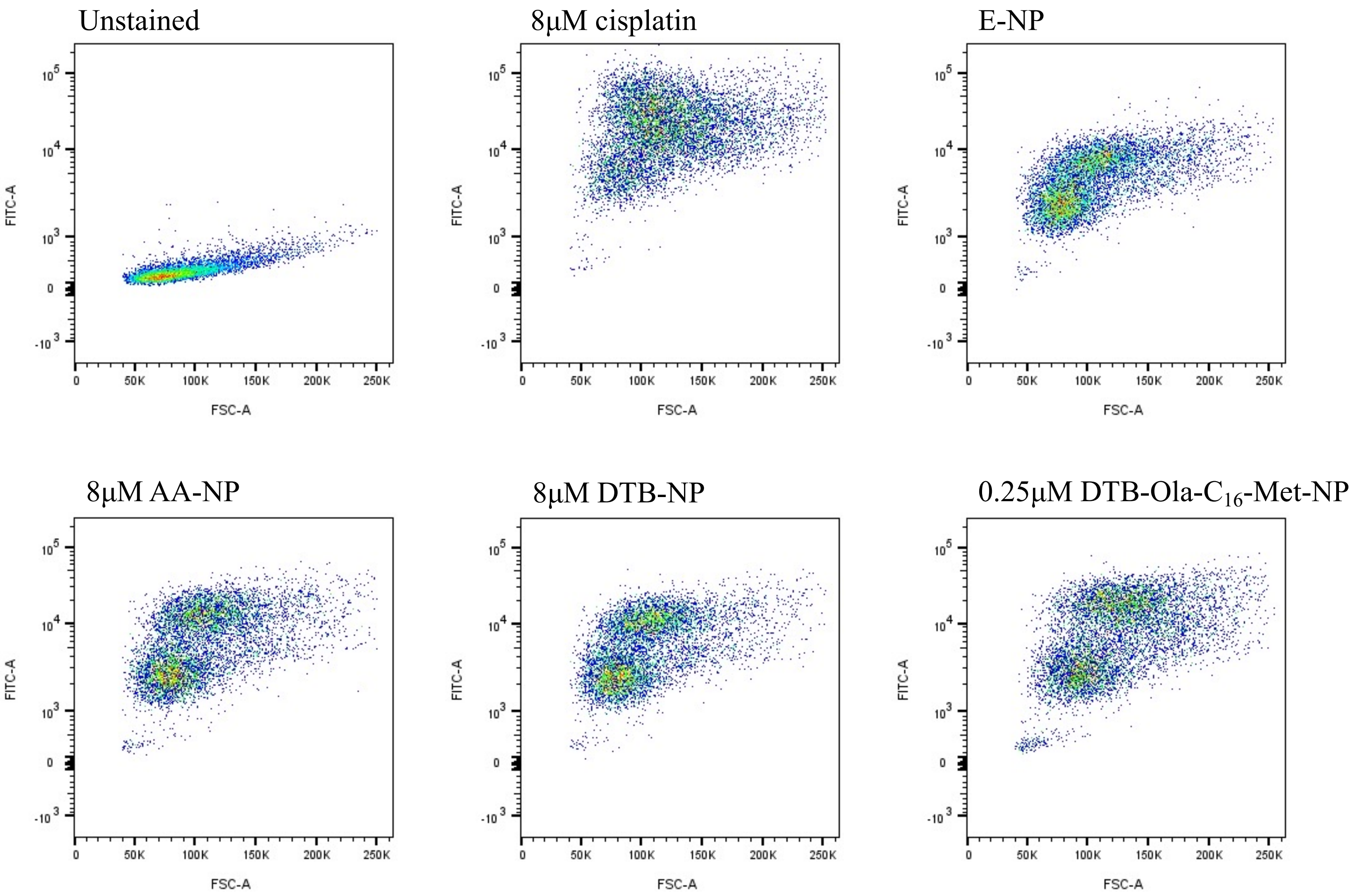


Fig. 6 from MS

### Western Blot of $C_{16}$-Met

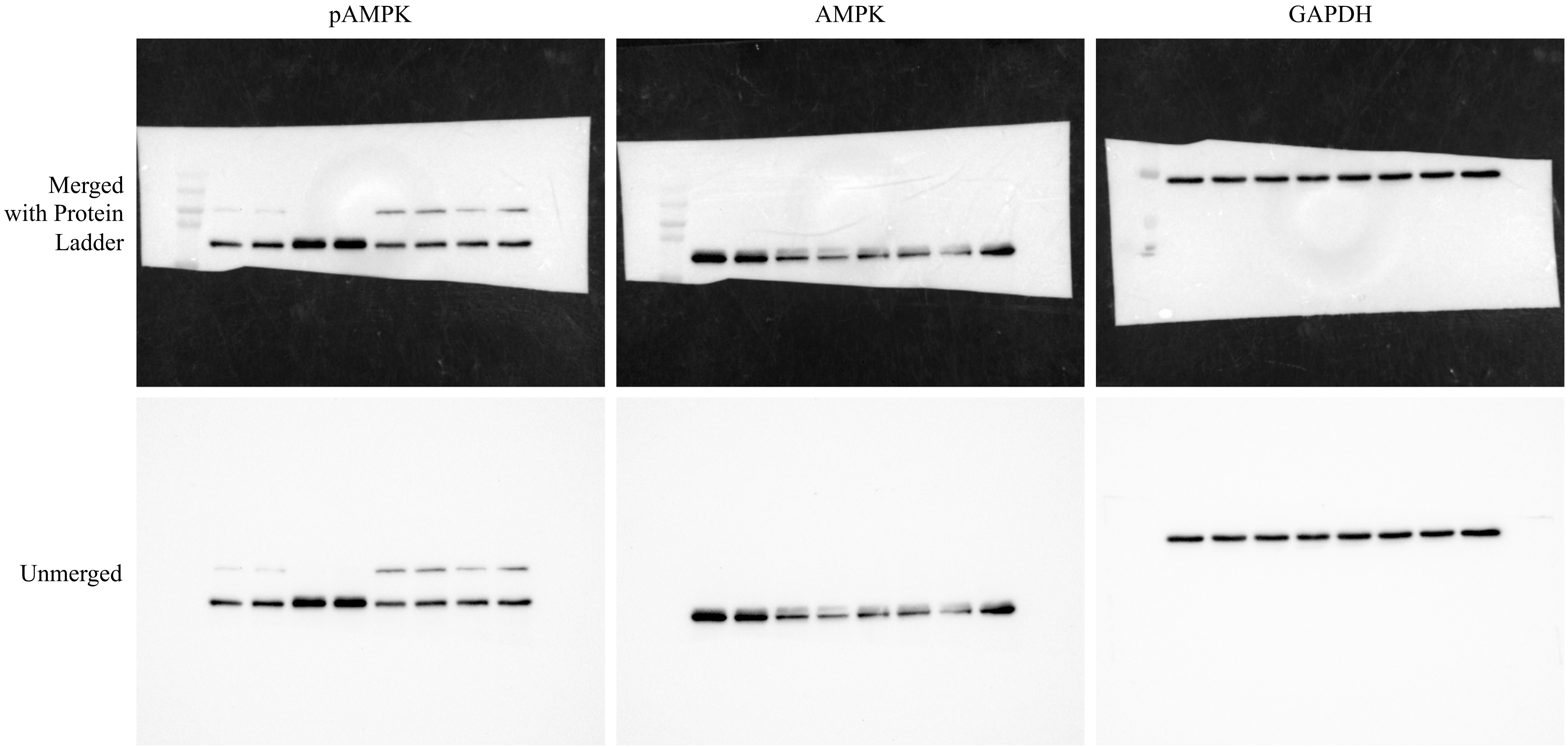


Fig. 6 from MS

### Western Blot of $C_6$-Met

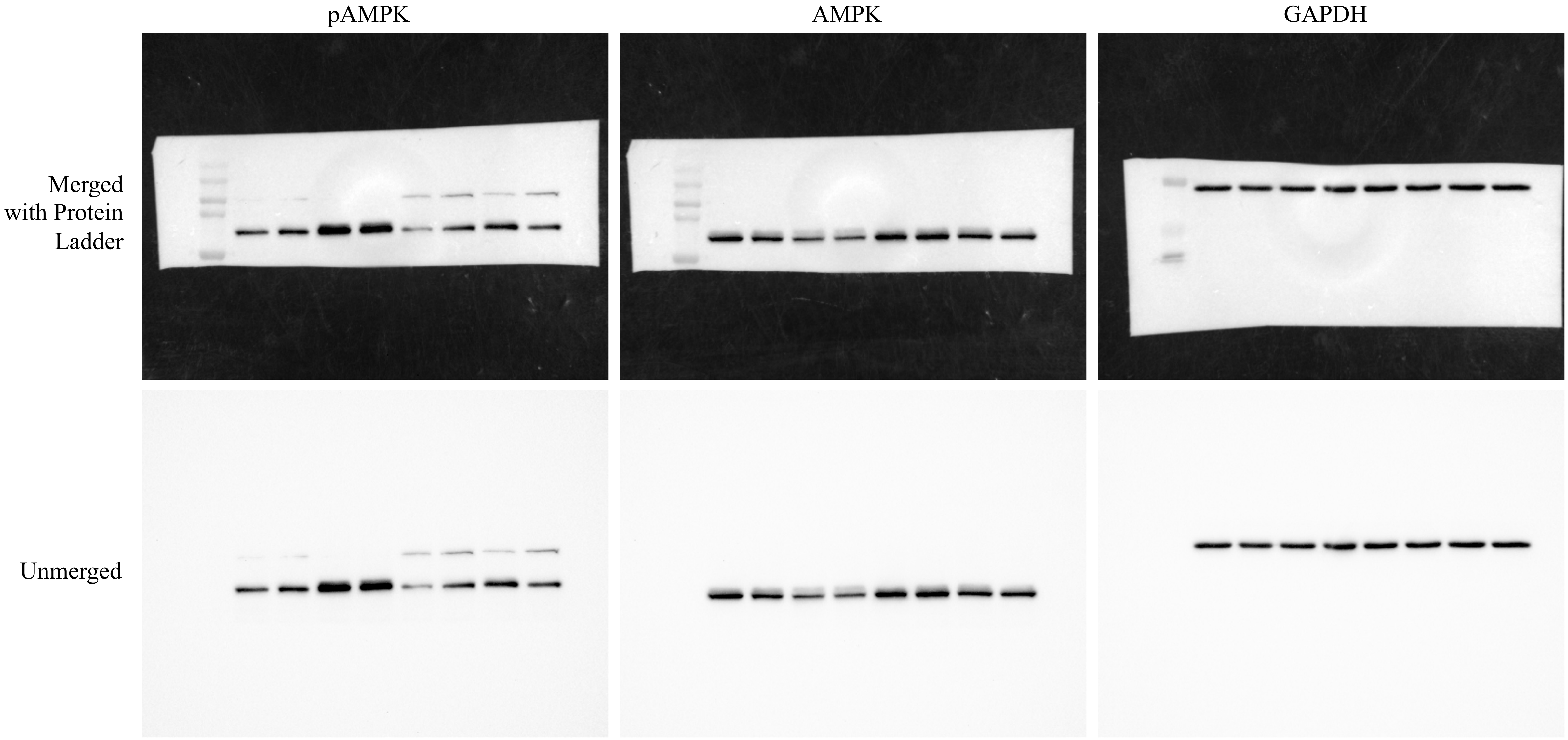


Fig. 6 from MS

### Western Blot of Ola NPs

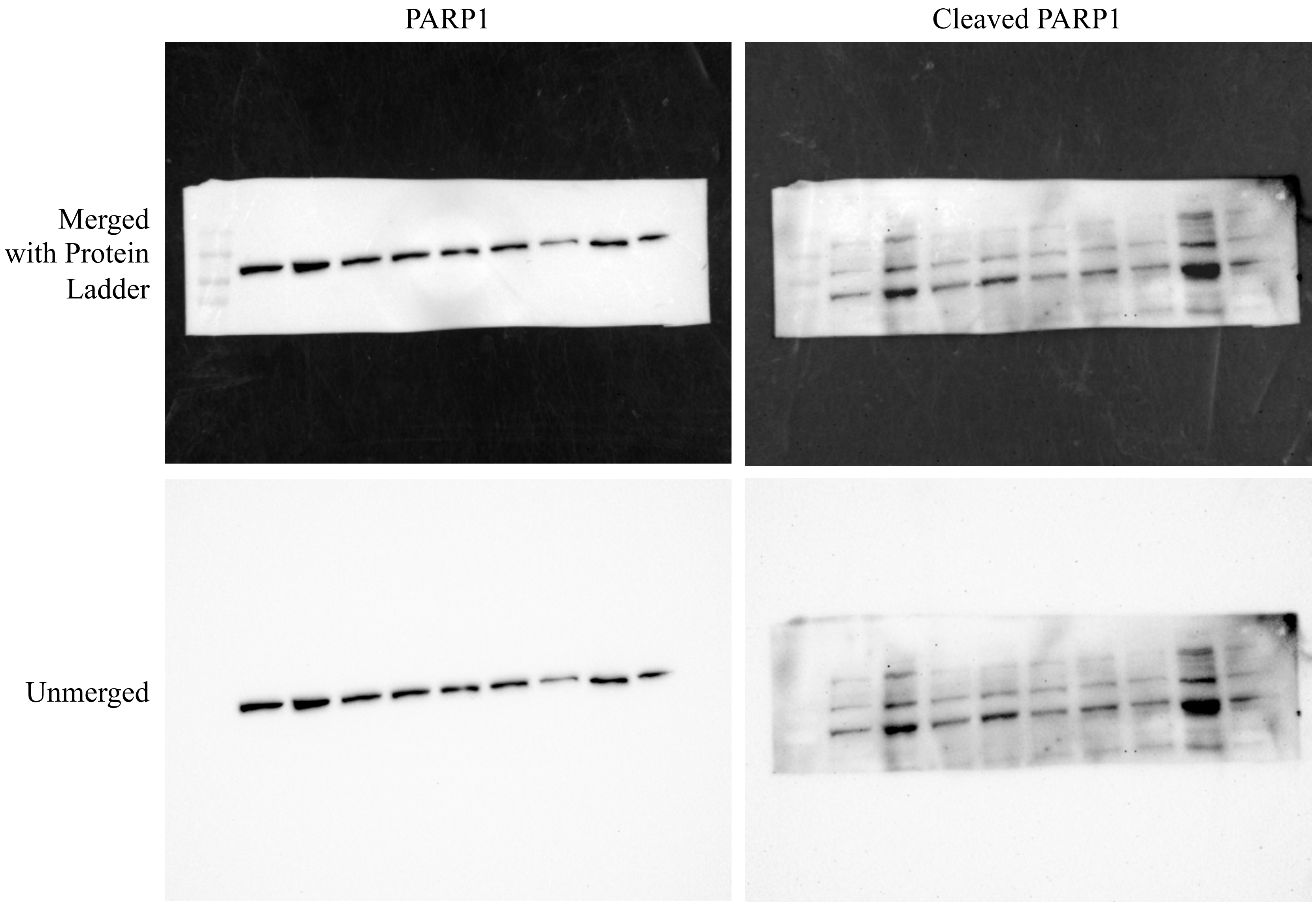


Fig. 6 from MS

#### Western Blot of Ola NPs

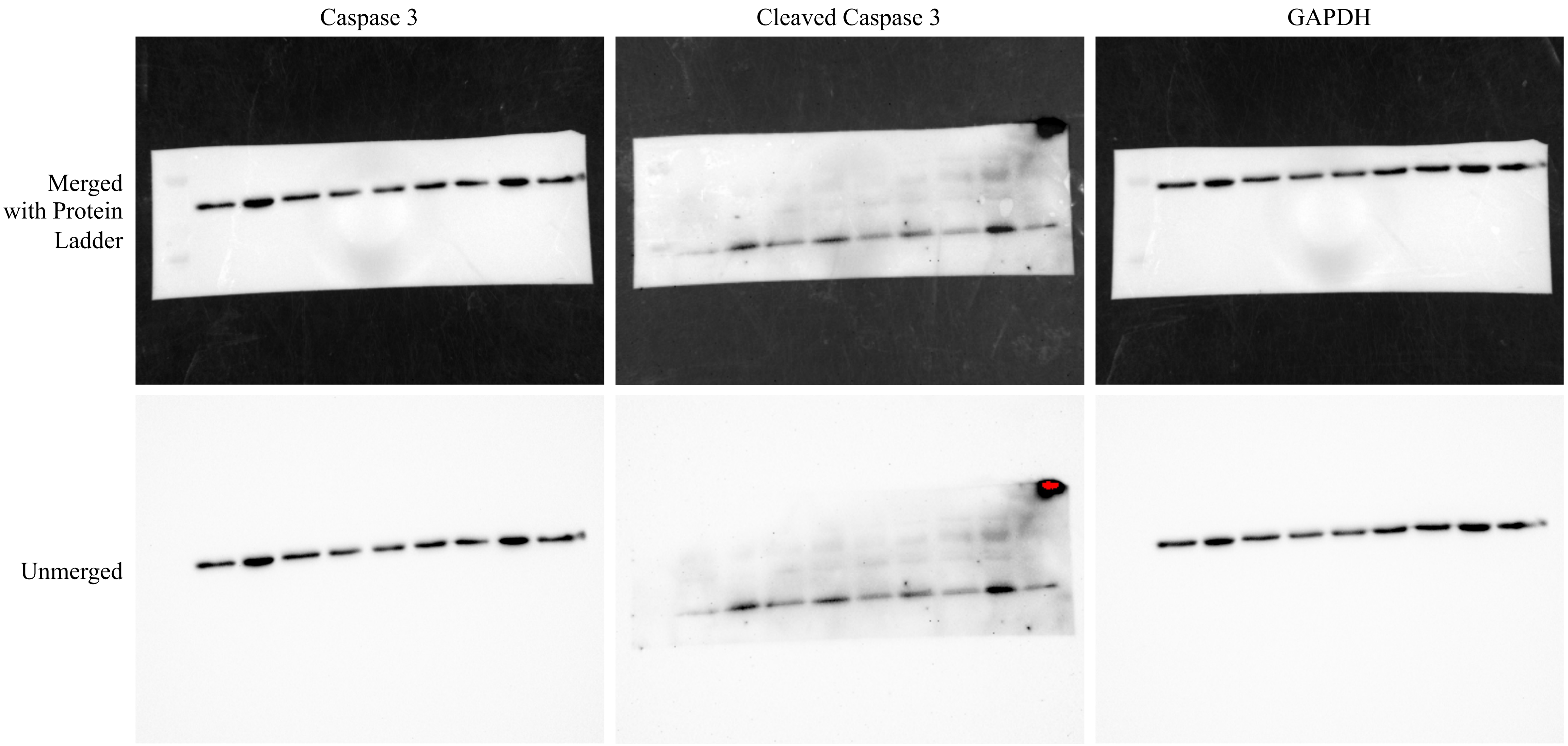


Fig. 6 from MS
